# A TET1-PSPC1-*Neat1* molecular axis modulates PRC2 functions in controlling stem cell bivalency

**DOI:** 10.1101/2021.08.25.457543

**Authors:** Xin Huang, Nazym Bashkenova, Yantao Hong, Diana Guallar, Zhe Hu, Vikas Malik, Dan Li, Xiaohua Shen, Hongwei Zhou, Jianlong Wang

## Abstract

TET1 maintains hypomethylation at bivalent promoters through its catalytic activity in embryonic stem cells (ESCs). However, whether and how TET1 exerts catalytic activity-independent functions in regulating bivalent genes is not well understood. Using a proteomics approach, we mapped the TET1 interactome in mouse ESCs and identified PSPC1 as a novel TET1 partner. Genome-wide location analysis reveals that PSPC1 functionally associates with TET1 and Polycomb repressive complex-2 (PRC2) complex. We establish that PSPC1 and TET1 repress, and *Neat1*, the PSPC1 cognate lncRNA, activates the bivalent gene expression. In ESCs, *Neat1* tethers the TET1-PSPC1 pair with PRC2 at bivalent promoters. During the ESC-to-formative epiblast-like stem cell (EpiLC) transition, PSPC1 and TET1 promote PRC2 chromatin occupancy at bivalent gene promoters while restricting *Neat1* functions in facilitating PRC2 binding to bivalent gene transcripts. Our study uncovers a novel TET1-PSPC1-*Neat1* molecular axis that modulates PRC2 binding affinity to chromatin and bivalent gene transcripts in controlling stem cell bivalency.

**In Brief:** TET1 is a transcriptional repressor for bivalent genes in pluripotent stem cells, but its mechanistic action on stem cell bivalency is unclear. Huang et al. use proteomics and genetic approaches to reveal that catalytic activity-independent functions of TET1, coordinated with the paraspeckle components PSPC1 and its cognate lncRNA *Neat1*, dynamically regulates stem cell bivalency by modulating PRC2 binding affinity to chromatin and bivalent gene transcripts in pluripotent state transition.

**Highlights:**

- The TET1 interactome identifies PSPC1 as a novel partner in ESCs
- TET1 and PSPC1 repress bivalent genes by promoting PRC2 chromatin occupancy
- *Neat1* facilitates bivalent gene activation by promoting PRC2 binding to their mRNAs
- *Neat1* bridges the TET1-PSPC1 and PRC2 complexes in regulating bivalent gene transcription

## INTRODUCTION

Embryonic stem cells (ESCs) and epiblast stem cells (EpiSCs) represent the naïve and primed pluripotency states, respectively. They differ significantly in their epigenomic and transcriptomic features, clonogenicity, and differentiation potentials (Nichols and Smith, 2009). Accumulating evidence suggests that mammalian epiblast development may possess a series of intermediate pluripotent states in the developmental continuum between the naïve and primed phase, including formative pluripotency (Morgani et al., 2017; Smith, 2017). Epiblast-like stem cells (EpiLCs), a kind of formative pluripotent cells, transiently emerge when adapting ESCs to primed EpiSC culture conditions within a specific period (usually 48 hours), while an extended culture of EpiLCs establishes a stable primed state (Hayashi et al., 2011). Recently, stable cell lines with features of a formative state were generated by further modifying the culture conditions with specific combinations of cytokines and inhibitors (Kinoshita et al., 2021; Wang et al., 2021; Yu et al., 2021). Notably, a unique molecular feature of formative pluripotency, i.e., the “super-bivalency” at lineage-specific genes, was recently revealed from both *in vivo* E6.5 epiblast (Xiang et al., 2020) and *in vitro* formative cell lines (Wang et al., 2021). In mammals, gene promoters marked by both H3K4me3 and H3K27me3 are termed bivalent promoters. These bivalent domains are considered to poise the expression of developmental regulators in ESCs while allowing timely activation upon differentiation cues (Voigt et al., 2013). KMT2B (MLL2) and Polycomb Repressive Complex-2 (PRC2) are responsible for depositing H3K4me3 and H3K27me3, respectively, at bivalent promoters in ESCs (Boyer et al., 2006; Hu et al., 2013). DNA methylation at bivalent promoters decreases KMT2B activity and H3K4me3, and the loss of H3K4me3 leads to increased PRC2 occupancy at promoters (Mas et al., 2018). TET (ten-eleven translocation) family of proteins epigenetically regulate gene expression through DNA demethylation, converting 5-methylcytosine (5mC) to 5-hydroxymethylcytosine (5hmC) and other oxidized derivatives (Kohli and Zhang, 2013). It was thus suggested that TET proteins might play a pivot role in regulating bivalency (Mas et al., 2018; Xiang et al., 2020).

The TET family of proteins is expressed in various tissues and cell types. While the loss of TET proteins (*Tet1*KO or *Tet1/2/3*TKO) causes global changes in the DNA epigenome and gene expression in both mouse and human ESCs, the cells nevertheless retain the ability to self-renew (Dawlaty et al., 2011; Lu et al., 2014; Verma et al., 2018). In the formative EpiLCs and the primed EpiSCs, TET1 is the only expressed TET protein (Fidalgo et al., 2016; Khoueiry et al., 2017). Loss of TET1 caused dysregulation of gene expression in multiple ESC differentiation models (Dawlaty et al., 2011; Koh et al., 2011) and defects in mouse post-implantation development (Khoueiry et al., 2017). Mechanically, TET1 activates and represses gene transcription by catalytic activity-dependent functions in promoter/enhancer demethylation (Kohli and Zhang, 2013) and catalytic activity-independent functions in association with the epigenetic repressors, including the SIN3A/HDAC (Williams et al., 2011) and PRC2 (Neri et al., 2013; Wu et al., 2011) complexes, respectively. In addition, TET1 is also responsible for maintaining the DNA methylation valleys at promoters of developmentally regulated genes to establish a super-bivalency in the post-implantation epiblast (Xiang et al., 2020). However, whether and how the catalytic activity-independent functions of TET1 may also play a role in regulating bivalent genes have not been defined.

PRC2 modifies the chromatin to maintain the developmental lineage genes in their repressive state in ESCs (Boyer et al., 2006; Hojfeldt et al., 2019). Interestingly, TET1 was reported to repress the expression of bivalent genes through PRC2 recruitment, although the direct interaction between TET1 and PRC2 could not be demonstrated (Wu et al., 2011). The post-transcriptional gene regulation by PRC2 has been increasingly appreciated through discovering its association with RNA-binding proteins (RBPs) and long noncoding RNA (lncRNAs) that can regulate gene expression *in cis* or *in trans* (Cifuentes-Rojas et al., 2014; Davidovich and Cech, 2015; Kaneko et al., 2014; Yan et al., 2019). In addition to the functional association of lncRNAs with PRC2, nascent mRNAs and other RNA transcripts were also proposed to antagonize the association of PRC2 with the chromatin (Beltran et al., 2016; Davidovich et al., 2015; Kaneko et al., 2013; Long et al., 2020; Wang et al., 2017b). *In vivo*, a “PRC2 eviction” model was proposed in which the nascent mRNA regulates its own production by evicting PRC2 from the promoter, thereby further promoting gene transcription (Skalska et al., 2021; Wang et al., 2017a). Although TET1 could also be an RBP (He et al., 2016), whether TET1 may functionally connect with PRC2 through other RBPs and/or lncRNAs to control bivalent genes in pluripotent states has not been determined.

Here, through the study of TET1-associated proteins in mouse ESCs, we report the discovery of Paraspeckle component 1 (PSPC1), an RBP generally associated with nuclear paraspeckles (Knott et al., 2016), as a novel TET1 partner. We further establish that PSPC1 and its cognate lncRNA *Neat1* associate with TET1 and PRC2 at bivalent promoters. Using genetic loss-of-function approaches, we demonstrate that TET1 and PSPC1 tether PRC2 on the chromatin through *Neat1* to inhibit the PRC2 binding to bivalent gene transcripts, thereby preventing PRC2 eviction from chromatin during pluripotent state transition. On the other hand, upon the loss of TET1 or PSPC1, *Neat1* enhances PRC2 affinity to mRNAs, thereby activating transcription of bivalent genes. Our study thus establishes a previously unappreciated TET1-PSPC1-*Neat1* molecular axis that modulates PRC2 affinity to chromatin and bivalent gene transcripts in controlling stem cell bivalency.

## RESULTS

### The TET1 interactome in ESCs identifies PSPC1 as its interacting partner

We engineered mouse ESCs expressing FLAG-tagged *Tet1* (FL-*Tet1*) and purified the TET1 protein complexes using SILAC (stable isotope labeling by amino acid in cell culture)-based AP-MS (affinity purification followed by mass spectrometry) method as described in our previous studies (Ding et al., 2015; Guallar et al., 2018; Huang et al., 2021). Reciprocal SILAC labeling was performed as biological replicates, and the protein intensity ratios of FLAG (TET1) versus Control immunoprecipitation (IP) (Rep1: light/heavy; Rep2: heavy/light) for each protein were plotted (Figure 1A; Table S1). Validating our approach, we identified several known TET1 partners such as OGT and SIN3A (Vella et al., 2013) and components of a ribosome biogenesis complex consisting of PELP1, TEX10, WDR18, and SENP3 (Finkbeiner et al., 2011), consistent with our previous finding that TET1 and TEX10 are close partners (Ding et al., 2015). In addition, we identified several RNA binding proteins (RBPs), such as L1TD1 and PSPC1 (Figures 1A and S1A). Selected candidate proteins in the TET1 interactome were validated by FLAG co-immunoprecipitation (co-IP) followed by western blot (WB) analysis (Figure 1B). We decided to focus on the TET1 and PSPC1 partnership for several reasons. First, while the functional significance of the TET1-SIN3A/OGT (Deplus et al., 2013; Vella et al., 2013; Williams et al., 2011) and TET1-TEX10 (Ding et al., 2015) partnerships in ESC maintenance or differentiation is well-established, the functional cooperation between TET1 and PSPC1 is unclear. PSPC1 does interact with other paraspeckle components such as SFPQ and NONO in ESCs (Figure S1B), although paraspeckles were not observed in mouse (Figure S1C) or human (Chen and Carmichael, 2009) ESCs. Second, TET1 activates and represses lineage gene expression during ESC differentiation through its catalytic activity-dependent and -independent functions, respectively (Koh et al., 2011; Wu et al., 2011; Zhu et al., 2018). While TET1 catalytic activity-independent function may act through PRC2, their direct physical association was not detected (Wu et al., 2011), raising the possibility of unknown bridging proteins and/or RNAs for the functional interaction between TET1 and PRC2.

**Figure 1.**
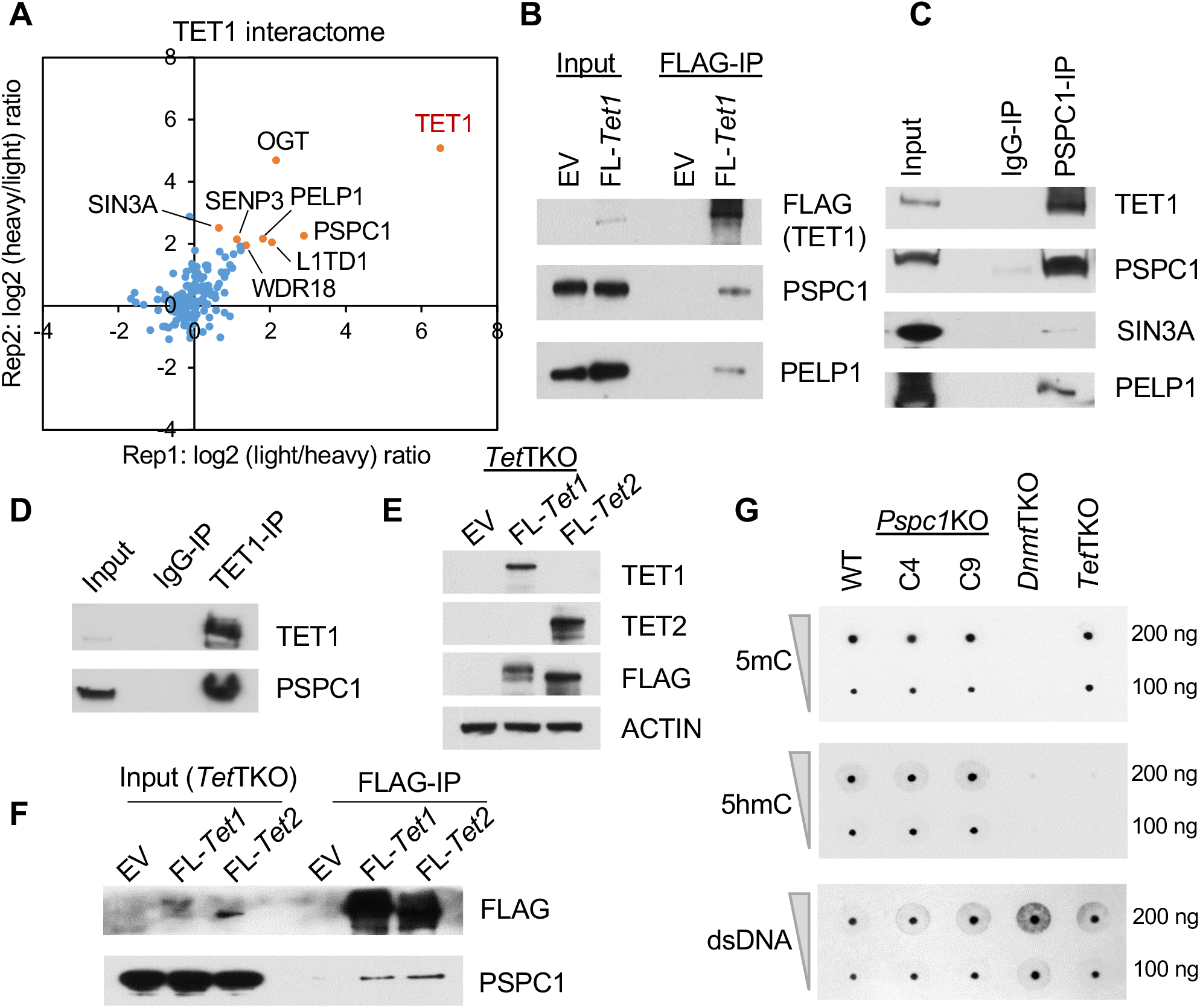
PSPC1 is a novel partner of TET1 in ESCs. (A) Protein ratios of FLAG-IP (TET1) versus Control-IP (empty vector) AP-MS in two replicates with reciprocal SILAC labeling are plotted, and a few proteins in the TET1 interactome are indicated. (B) Co-immunoprecipitation (co-IP) of TET1 partners by FLAG-IP followed by western blot analysis with indicated antibodies. (C-D) co-IP by endogenous PSPC1 (C) and TET1 (D) antibodies followed by western blot analysis with indicated antibodies. (E) *Tet1/2/3* triple KO (*Tet*TKO) ESCs are rescued with FLAG-tagged TET1 or TET2. (F) Both TET1 and TET2 interact with PSPC1. FLAG-IP of TET proteins and western blot analysis of PSPC1 are presented. (G) DNA 5mC and 5hmC dot blot analysis of WT and *Pspc1*KO (two independent clones C4 and C9) ESCs. dsDNA antibody is reblotted as the loading control. *Dnmt1/3a/3b* triple KO (*Dnmt*TKO) and *Tet*TKO ESCs serve as negative controls of 5mC and 5hmC, respectively.

We confirmed that PSPC1 and TET1 interact by reciprocal co-IP using endogenous antibodies (Figure 1C-D). PSPC1 also interacts with the TET1 partners such as SIN3A and PELP1, although with a much weaker affinity than TET1 (Figure 1C). We previously reported that PSPC1 also interacts with TET2 in ESCs (Guallar et al., 2018). To probe the potential biochemical entities associated with PSPC1, TET1, and TET2, we performed size exclusion chromatography (i.e., gel filtration) experiments on ESC nuclear extracts. We found the co-fractionation of all these three factors (Complex I, blue, Figure 1E) as well as the TET1-free co-fractionation of PSPC1 and TET2 (Complex II, red, Figure S1E). While the existent TET1-free TET2/PSPC2 complex has been demonstrated with the unique function of TET2, but not TET1, in RNA-dependent chromatin targeting for ERV control in ESCs (Guallar et al., 2018), here we decided to address whether the PSPC1/TET1 interaction is mediated by TET2 in light of their co-fractionation as Complex I in ESC nuclear extracts (Figure S1E). We thus employed the *Tet1/2/3* triple KO (*Tet*TKO) ESCs (Fidalgo et al., 2016), rescued with either FLAG-tagged TET1 or TET2 (Figure 1E, TET3 is not expressed in ESCs), and performed FLAG-IP followed by WB analysis of PSPC1. Interestingly, we found that both TET1 and TET2 interact with PSPC1 in the absence of the other TET proteins (Figure 1F), indicating the TET2-independent TET1-PSPC1 interaction and further confirming the TET1-independent TET2-PSPC1 interaction, despite the co-fractionation of these three proteins in size exclusion chromatography.

We also performed domain mapping experiments to dissect the TET1/PSPC1 interaction. The full-length (2,039 amino acids) or truncated fragments of *Tet1* were cloned into the FLAG-tagged expression vectors (Figure S1F) for transfection in ESCs followed by Co-IP. We observed that full-length *Tet1* and its variants (C1, C2, and ΔCXXC) containing a minimal C-terminal catalytic domain (amino acids 1,367∼2,039) interact with PSPC1 (Figure S1F). Similarly, we cloned the full-length or truncated fragments of *Pspc1* into the V5-tagged expression vectors (Figure S1G) for transfection in ESCs followed by Co-IP. We found that full-length PSPC1 and its truncated variant F2 containing the multifunctional Drosophila behavior/human splicing (DBHS) domain (Knott et al., 2016) were required to interact with TET1 (Figure S1G). We then asked whether PSPC1 participates in the catalytic activity-dependent functions of TET1 in ESCs. We employed *Pspc1*KO ESCs (two independent clones, shown in Figure S1D) (Guallar et al., 2018) and performed DNA dot-blot analysis. We found that PSPC1 ablation does not affect the DNA 5mC or 5hmC intensity in ESCs (Figure 1G).

Taken together, we identified PSPC1 as a novel TET1 partner that may modulate TET1 functions in ESC pluripotency independently of its catalytic activity.

### PSPC1, TET1, and PRC2 co-localize at the bivalent gene promoters in ESCs

PSPC1 is a DNA- and RNA-binding protein (Knott et al., 2016). To understand the function of PSPC1 in pluripotency, we performed chromatin-immunoprecipitation followed by deep sequencing (ChIP-seq) analysis of PSPC1 in WT and *Pspc1*KO ESCs. We identified 2,324 PSPC1 ChIP-seq peaks in ESCs, using PSPC1 ChIP in *Pspc1KO* cells as the background. The majority (74.2%) of PSPC1 binding peaks are located at the gene promoters, within 5K bp of transcriptional start sites (TSSs), with PSPC1 ChIP signal also enriched at TSSs (Figure 2A-B). Consistent with the PSPC1-TET1 partnership, almost all PSPC1 peaks (91.7%, 2,132/2,324) co-localize with TET1 binding regions (Figure S2A). We compared the DNA 5hmC and 5mC intensities at the TET1 peak regions with or without PSPC1 occupancy from published (hydroxy)methylated DNA immunoprecipitation sequencing (hme/meDIP-seq) data in ESCs (Xiong et al., 2016). Overall, the PSPC1/TET1 common regions lack 5hmC and 5mC compared with the TET1-only regions (Figure S2B), consistent with our finding that PSPC1 may not participate in the catalytic activity-dependent functions of TET1 in ESCs (Figure 1G).

**Figure 2.**
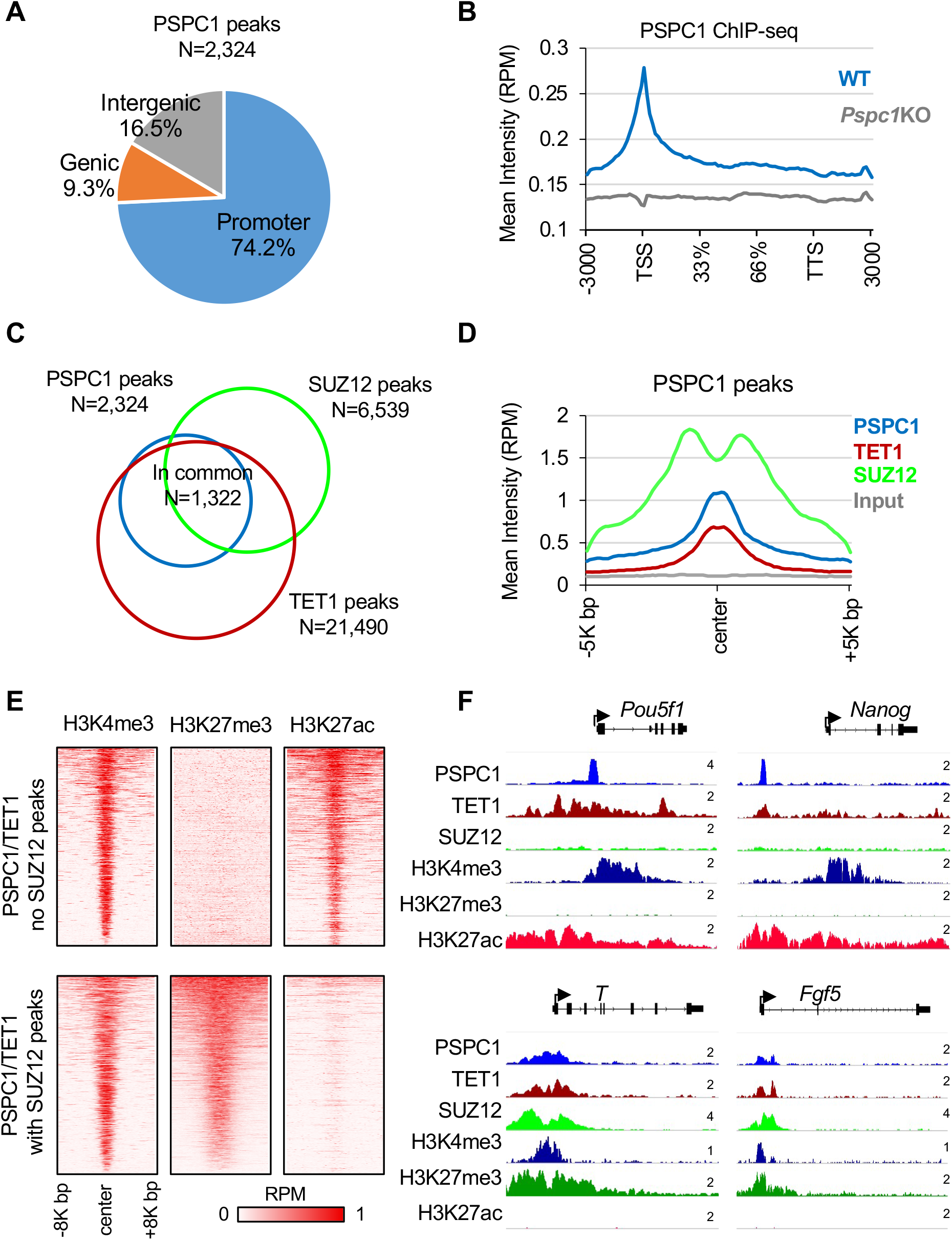
PSPC1, TET1, and PRC2 co-localize at bivalent promoters in ESCs. (A) Annotation of PSPC1 ChIP-seq peaks in ESCs at promoters (distance to TSS < 5K bp), intergenic or genic regions. (B) Mean intensity plot by reads per million (RPM) shows PSPC1 ChIP-seq intensity of WT and *Pspc1*KO ESCs at gene bodies (within ± 3000 bp). TSS, transcription start site, TTS, transcription termination site. (C) Overlap of the PSPC1, TET1, and PRC2 subunit SUZ12 peaks in ESCs. TET1 ChIP-seq data in ESCs are from (Wu et al., 2011). (D) Mean intensity plot by RPM shows PSPC1, TET1, and RPC2 subunit SUZ12 ChIP-seq intensity at PSPC1 peak regions (within ± 5K bp around PSPC1 peak center). (E) Heatmaps by RPM shows histone marks H3K4me3 and H3K27ac (Hon et al., 2014) and H3K27me3 (Cruz-Molina et al., 2017) at PSPC1/TET1 common peak regions (within ± 8K bp around PSPC1 peak center) with and without PRC2 occupancy. (F) ChIP-seq tracks of PSPC1, TET1, SUZ12, and histone marks of H3K4me3, H3K27me3, H3K27ac at PSPC1/TET1 common peak regions with (*T, Fgf5*) and without (*Pou5f1, Nanog*) PRC2 occupancy. The numbers indicate the normalized RPM value of the tracks shown.

To understand how PSPC1 may functionally interact with other pluripotency-related transcription factors or epigenetic regulators, we performed the ChIP-seq correlation analysis (Ding et al., 2015) to examine their genome-wide binding patterns. We found that PSPC1 DNA binding sites are more like those of TET1 and EZH2/SUZ12 (Figure S2C), suggesting that PSPC1 may be involved in TET1- and PRC2-dependent regulations. Indeed, 56.9% (1,322/2,324) of the PSPC1 peaks are co-occupied by TET1 and PRC2 component SUZ12 (Figure 2C). TET1 and SUZ12 are also enriched at PSPC1-bound regions (Figure 2D). PRC2 deposits the repressive histone mark H3K27me3, coexistent with the active histone mark H3K4me3 at the promoters of bivalent genes in ESCs that are lowly expressed and poised to be promptly activated upon differentiation (Boyer et al., 2006). Consistently, gene ontology (GO) analysis for the PSPC1/TET1/SUZ12 common targets revealed that many of the genes are involved in organism development, cell fate commitment, and cell differentiation (Figure S2D). Next, we compared the intensity of histone marks H3K4me3, H3K27ac, and H3K27me3 at the PSPC1/TET1 common peaks with or without SUZ12 occupancy. The PSPC1/TET1 peaks without SUZ12 occupancy are enriched with active marks of H3K4me3 and H3K27ac (e.g., the promoters of *Pou5f1* and *Nanog*), whereas the PSPC1/TET1/SUZ12 common peaks are enriched with bivalent marks of H3K4me3 and H3K27me3 (e.g., the promoters of *T* and *Fgf5*) (Figure 2E-F).

Together, these results suggest a potential physical association of the TET1-PSPC1 partnership with PRC2 in repressing bivalent genes in ESCs. However, the possible role of the TET1-PSPC1 partnership independent of PRC2 in activating pluripotency genes cannot be discounted and warrants future investigation (see Discussion).

### PSPC1 restricts bivalent gene activation during the ESC-to-EpiLC transition

To understand how PSPC1 may contribute to the regulation of bivalent genes in pluripotent cells, we decided to study the functions of PSPC1 in the pluripotent state transition, during which the super-bivalency of a large set of developmental genes was initially proposed (Morgani et al., 2017; Smith, 2017) and subsequently confirmed (Wang et al., 2021) in formative pluripotent stem cells. *Pspc1*KO does not affect the maintenance of ESCs (Guallar et al., 2018), making the ESC-to-EpiLC transition model applicable for such functional studies. By switching the culture medium from serum and LIF (SL) to Fgf2 and Activin A (FA), ESCs enter a transient formative pluripotency state of EpiLCs, followed by a primed pluripotency state of EpiSCs under an extended culture of EpiLCs in the FA condition (Smith, 2017). We thus adapted WT and *Pspc1*KO ESCs (D0) in FA culture medium for 2 days (D2) and 4 days (D4) and collected RNAs for RNA-seq analysis (Figure 3A). Of note, the D2 EpiLCs are considered as the state of formative pluripotency (Buecker et al., 2014; Fidalgo et al., 2016; Hayashi et al., 2011), while D4 EpiLCs and EpiSCs are of primed pluripotency when the meso/ectodermal lineage genes (e.g., *Fgf5, Fgf8, T, Eomes*, and *Otx2*) are further activated (Huang et al., 2017). Principal component analysis (PCA) revealed a trajectory of gene expression profiles moving from D0 (ESC) to D2 and D4 (EpiLC) on PC1, while the differences of gene expression between WT and KO cells at all 3 time points are reflected on PC2 (Figure 3B). By comparing the differentially expressed genes (DEGs, P-value<0.05, fold-change>1.5, Table S2) between WT and *Pspc1*KO cells at 3 time points, we found that multiple signaling pathways and their associated genes, including FGF signaling (e.g., *Fgf5* and *Fgf8*), Nodal signaling (e.g., *Nodal* and *Eomes*), and Wnt signaling (e.g., *Axin2, Wnt5b*, and *Wnt8a*), are upregulated in *Pspc1*KO relative to WT EpiLCs (D2 and D4, Figure S3A). GO analysis of these PSPC1-repressed DEGs in D2 and D4 EpiLCs indicates that they are involved in embryo and tissue development (Figure S3B). In contrast, the PSPC1-activated DEGs are involved in multiple cellular regulations, including metabolic process, protein transport, and cell death (Figure S3B). Interestingly, a majority (75.9%, 129/170) of the PSPC1-repressed DEGs in EpiLCs are not repressed by PSPC1 in ESCs (Figure S3B, left panel), likely due to their low expression levels and/or alternative repression mechanisms in ESCs.

**Figure 3.**
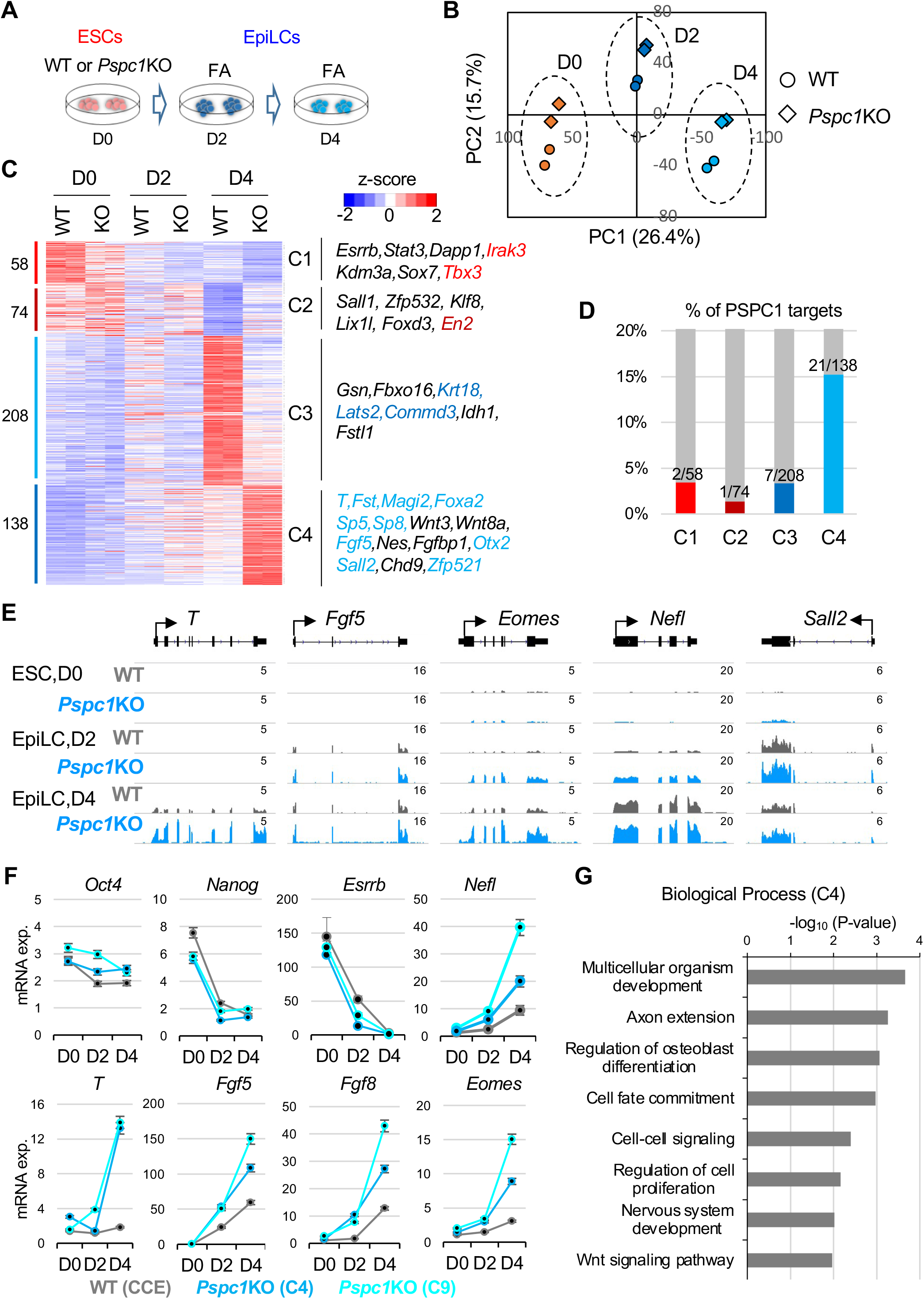
PSPC1 negatively regulates activation of bivalent genes in the pluripotent state transition. (A) Schematic depiction of the naïve-to-formative transition of WT and *Pspc1*KO ESCs. The ESCs are adapted in Fgf2 and Activin A (FA) culture medium for 2 days and 4 days. The ESCs (D0) and formative EpiLCs (D2, D4) are collected for RNA-seq analysis. (B) Principal component analysis (PCA) of WT and *Pspc1*KO RNA-seq samples at different time points. Percentages of variance explained in each principal component (PC) are indicated. (C) Heatmap showing the relative expression of differentially expressed genes (DEGs, by P-value<0.05, fold-change>1.5) by comparing D0 WT and D4 WT cells and comparing D4 WT and D4 KO cells. The numbers of DEGs are shown on the left, and representative genes in the four classes (C1-C4) are listed on the right with direct PSPC1 targets from ChIP-seq analysis indicated by the color text. (D) Histogram shows the percentages (%) and numbers of DEGs in each class (C1-C4) as the PSPC1 ChIP-seq targets. (E) RNA-seq tracks of WT and *Pspc1*KO ESCs and EpiLCs at bivalent lineage gene loci. The numbers indicate the normalized reads per million (RPM) value of the tracks shown. (F) RT-qPCR analysis of pluripotency and lineage genes in WT and *Pspc1*KO ESCs (CCE background with two independent clones, C4 and C9) during ESC-to-EpiLC differentiation. Error bars represent the standard deviation of triplicates. (G) Gene ontology (GO) analysis for the C4 genes (class 4, N = 138) shown in panel (C).

Next, we focused on the DEGs between D0 and D4 (ESC vs. EpiLC) WT cells and between D4 WT and *Pspc1*KO EpiLCs to obtain 478 shared DEGs by both comparisons (Figure 3C; Table S2). Clustering analysis of these genes illustrated different expression patterns among the samples (class1-4 or C1-4, Figure 3C). We examined the number of DEGs in classes C1-4 that are direct PSPC1 targets from ChIP-seq analysis and found that C4 contains the highest percentage (15.2%, 21/138) of PSPC1 targets (Figure 3D). These PSPC1 targets (e.g., *T, Fgf5*, and *Sall2*) are bivalent and lowly expressed genes in ESCs while transcriptionally activated in WT EpiLCs, and PSPC1 depletion further increases their expression during EpiLC differentiation (Figures 3C and 3E-F). Consistent with the GO analysis on the PSPC1/TET1/PRC2 common targets (Figure S2D), GO analysis of these PSPC1-repressed C4 genes indicated that they are involved in multicellular organism development, cell fate commitment, and Wnt signaling pathways (Figure 3G). Of note, NONO is a close partner of PSPC1 (Figure S1B) that often functions together with PSPC1 (Knott et al., 2016), and NONO also interacts with TET1 in ESCs (Li et al., 2020). Like *Pspc1*KO, *Nono*KO is compatible with ESC maintenance (Ma et al., 2016) and causes upregulation of lineage genes (e.g., *T, Eomes*, and *Fgf8*) in EpiLCs (Figure S3C-D).

In sum, our results establish PSPC1 as a transcriptional repressor that restricts bivalent gene activation during the ESC-to-EpiLC transition.

### *Neat1* promotes bivalent gene activation during the ESC-to-EpiLC transition

PSPC1 as an RBP was well-known for its roles in binding lncRNA *Neat1*, which drives the formation of nuclear paraspeckles (Isobe et al., 2020; Nakagawa et al., 2011). However, pluripotent stem cells do not form paraspeckles, and thus the functional relationship between PSPC1 and *Neat1* in pluripotency is not fully understood. Neither is known whether *Neat1* plays any role in TET1 functions. Therefore, we designed two sgRNAs targeting the *Neat1* locus and performed CRISPR/Cas9 genome editing to delete the 6K bp region containing the short (*Neat1_1*) isoform of *Neat1* (Figure 4A), the only isoform expressed in ESCs (Isobe et al., 2020) and EpiLCs (Figure 4B). Of note, the long *Neat1_2* is a somatic isoform that functions in driving paraspeckle formation (Isobe et al., 2020) and is collaterally abrogated by our CRISPR deletion (Figure 4A). We thus collectively refer to *Neat1*KO hereafter. The *Neat1*KO ESCs maintain self-renewal, consistent with the dispensability of *Neat1* for mouse embryo development (Nakagawa et al., 2011). We adapted WT and *Neat1*KO ESCs in FA culture medium and collected RNAs at Day 0 (ESC), Days 2 and 4 (EpiLC) for RNA-seq analysis. We confirmed that only *Neat1_1* is expressed in ESCs and EpiLCs and its expression gradually decreases during EpiLC differentiation (Figure 4B, reduced signal strengths on the left panel and FPKM values on the right panel). By comparing the DEGs (P-value<0.05, fold-change>1.5, Table S3) between WT and *Neat1*KO cells at 3 time points, we observed many bivalent genes (e.g., *Fgf5, Fgf8, Wnt8a*, and *Nefl*) are downregulated in D2 or D4 EpiLCs upon *Neat1*KO compared to the WT cells (Figures 4E and S4A), an effect that is opposite to that of *Pspc1*KO in EpiLCs (Figures 3E-F and S3A). Fewer DEGs were identified in D4 EpiLCs relative to D0 ESCs and D2 EpiLCs upon *Neat1*KO (Figure S4A), likely due to the relatively low *Neat1* expression in D4 EpiLCs (Figure 4B).

**Figure 4.**
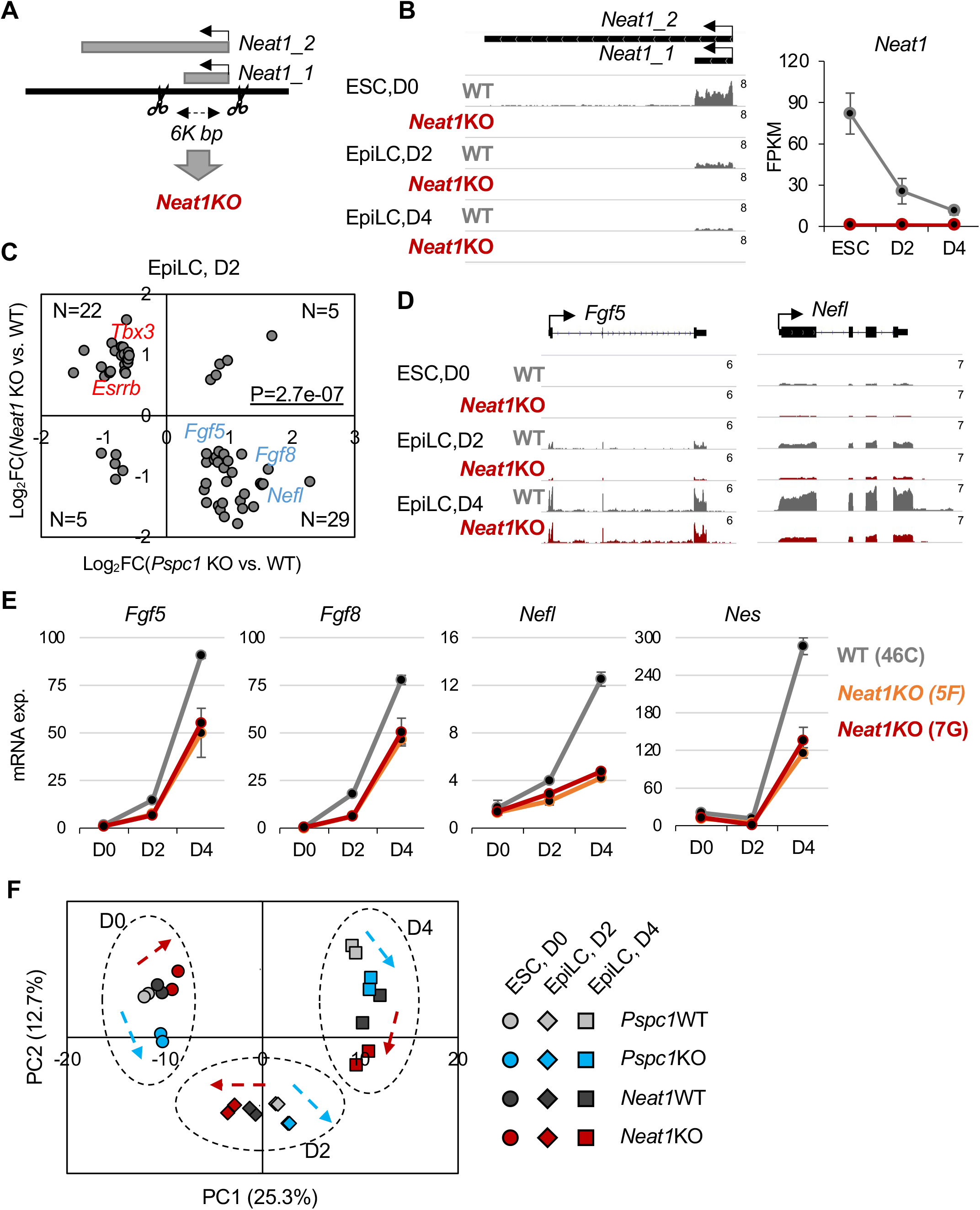
*Neat1* positively regulates bivalent gene activation in the pluripotent state transition. (A) Schematic depiction of the *Neat1*KO strategy. The scissors denote that gRNA targeting sites for CRISPR/Cas9 genome editing. The short (*Neat1_1*) and long (*Neat1_2*) isoforms of the mouse *Neat1* gene are indicated. (B) RNA-seq tracks (left) and expression of *Neat1* (right) during the ESC-to-EpiLC differentiation. The numbers indicate the normalized reads per million (RPM) value of the tracks shown (left). *Neat1* expression is shown in FPKM (fragments per kilobase of transcript per million mapped reads) values (right). Error bars represent the standard deviation of triplicates. (C) Scatter plot of the relative gene expression of DEGs (P-value<0.05, fold-change>1.5) upon the loss of *Pspc1* (*Pspc1* KO/WT) or *Neat1* (*Neat1* KO/WT) at D2 EpiLC from RNA-seq analysis. P-value was calculated using the Fisher-exact test. Representative genes are labeled on the plot. (D) RNA-seq tracks of WT and *Neat1*KO ESCs and EpiLCs at bivalent gene loci (*Fgf5* and *Nefl*). The numbers indicate the normalized reads per million (RPM) value of the tracks shown. (E) RT-qPCR analysis of bivalent genes in WT and *Neat1*KO ESCs (46C genetic background with two independent clones, 5F and 7G) during EpiLC differentiation. Error bars represent the standard deviation of triplicates. (F) Principal component analysis (PCA) of *Pspc1* WT and KO, *Neat1* WT and KO RNA-seq samples at different time points (D0, D2, D4). Percentages of variance explained in each principal component (PC) are indicated. The arrows indicate the trend of gene expression changes comparing the KO vs. WT samples at 3 time points.

To further investigate the functional relationship between PSPC1 and *Neat1*, we compared the RNA-seq gene expression ratios upon *Pspc1*KO and *Neat1*KO at 3 time points. We again observed a negative correlation of gene expression in ESCs (*r* = -0.27) and D2 EpiLCs (*r* = -0.23), but a weak positive correlation (*r* = 0.08) in D4 EpiLCs (Figure S4B). Next, we plotted the gene expression ratios of DEGs by *Pspc1*KO and *Neat1*KO at different time points. Interestingly, while *Pspc1*KO decreases and increases the expression of pluripotency (e.g., *Esrrb, Tbx3*) and bivalent (e.g., *Fgf5, Fgf8, Nefl*) genes, respectively, in D2 EpiLCs, as previously observed (Figures 3 and S3), *Neat1*KO exhibits an opposite effect in the regulation of those genes (Figures 4C-E and S4C). The PCA analysis of the *Pspc1*KO and *Neat1*KO RNA-seq samples shows that D0 (ESC) and D2 and D4 (EpiLC) samples group together and move towards right on PC1 during EpiLC differentiation (Figure 4F). Consistent with the correlation analysis (Figures 4C and S4B-C), the *Pspc1*KO and *Neat1*KO samples deviated to opposite directions compared with their WT samples in D0 and D2 (Figure 4F, refer to the direction of arrows at each time point).

Together, our results demonstrate that *Neat1* promotes bivalent gene activation during the ESC-to-EpiLC transition, establishing opposing functions of PSPC1 and its cognate lncRNA *Neat1* in controlling bivalent gene expression in pluripotent stem cells.

### PSPC1 and TET1, but not *Neat1*, are required for PRC2 chromatin occupancy at bivalent promoters during the ESC-to-EpiLC transition

We have established the co-occupancy of PSPC1 at the bivalent gene promoters in ESCs with TET1 and PRC2 (Figure 2C-F), two well-established repressors of lineage genes in ESCs and during differentiation (Khoueiry et al., 2017; Koh et al., 2011; Wu et al., 2011). To understand how PSPC1 and its cognate *Neat1* may modulate TET1 and PRC2 functions on transcriptional regulation of bivalent genes in pluripotent stem cells, we first asked whether PSPC1 contributes to TET1 and PRC2 chromatin binding. In ESCs, *Pspc1*KO doesn’t affect the chromatin-bound fraction of TET1 or PRC2 subunit SUZ12 (Figure S5A). SUZ12 ChIP-qPCR in ESCs also indicated that PRC2 binding at bivalent promoters is not affected by *Pspc1*KO (Figure S5B). We then addressed the potential roles of TET1 in the ESC-to-EpiLC transition. We established a degron system (Nabet et al., 2018) for rapid and inducible TET1 protein degradation with an extra benefit in knocking in an HA epitope tag to obviate the lack of a reliable TET1 antibody for downstream analyses (Figure 5A-B; see details in Methods). Using two independent *Tet1*-degron ESC lines, we confirmed that activation of lineage genes (e.g., *T, Fgf5, Fgf8*) during the ESC-to-EpiLC transition is further enhanced by dTAG13-treatment (i.e., TET1 depletion) (Figure S5C), phenocopying *Pspc1*KO (Figure 3F).

**Figure 5.**
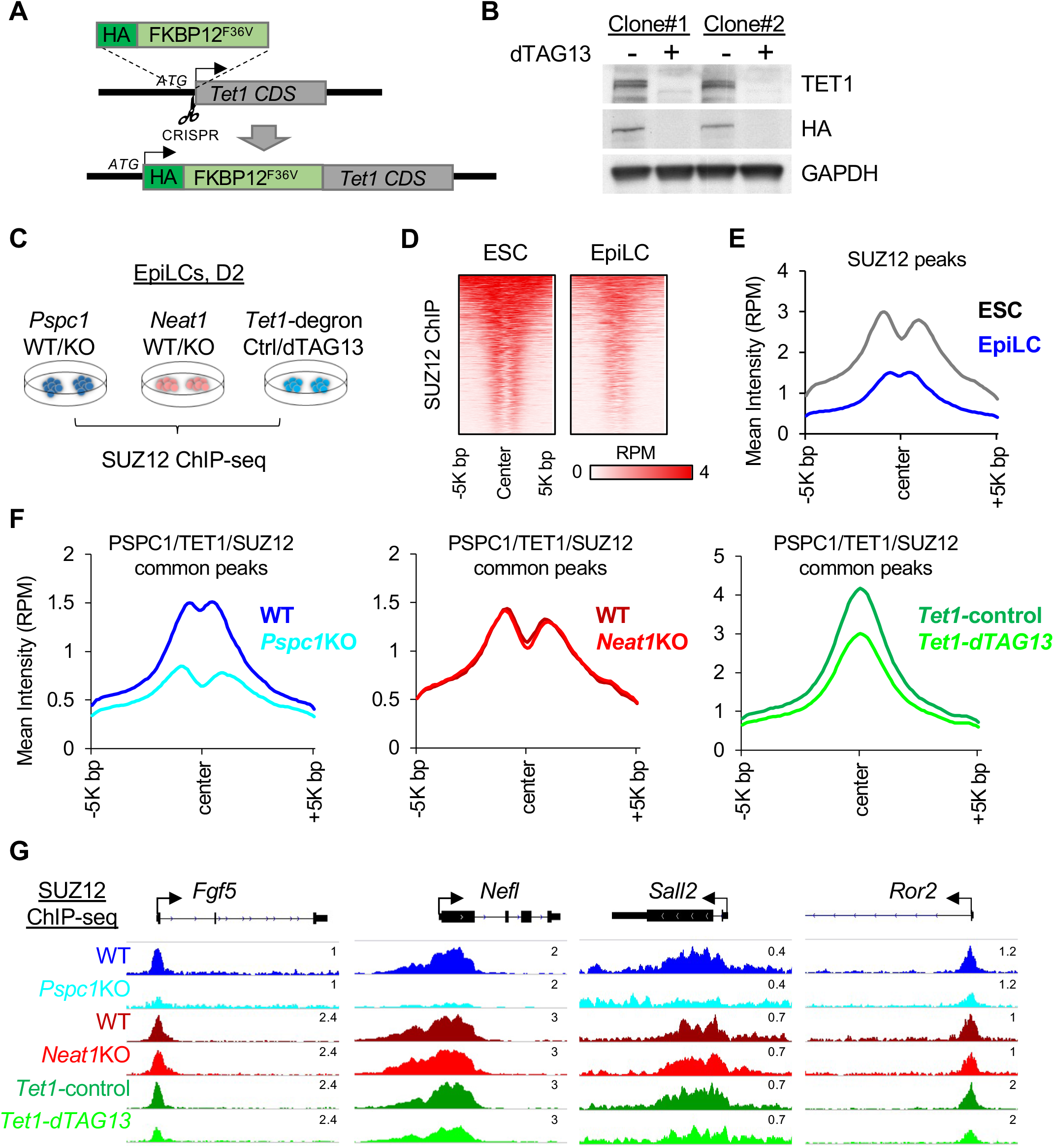
Depletion of PSPC1 or TET1 accelerates PRC2 eviction from bivalent promoters. (A) Schematic depiction of the *Tet1-*degron knock-in (KI) strategy using CRISPR/Cas9 genome editing (the Scissor symbol). The HA-tagged FKBP12^F36V^ donor sequence is inserted right after the start codon (ATG) of TET1 CDS to create the in-frame fusion protein. (B) Western blot analysis of TET1 protein in *Tet1*-degron ESCs (two independent clones) upon dTAG-13 treatment for 24 hrs. Degradation of TET1 was indicated by both endogenous antibody and HA fusion protein tag. (C) Schematic depiction of the PRC2 subunit SUZ12 ChIP-seq analysis in D2 EpiLCs of different genotypes (*Pspc1* WT/KO, *Neat1* WT/KO) or treatment (*Tet1*-degron with control/dTAG13). (D-E) Heatmap (D) and mean intensity plot (E) by reads per million (RPM) of SUZ12 ChIP-seq intensity at SUZ12 peak regions (within ± 5K bp around SUZ12 peak center, identified in ESCs) in WT ESCs and EpiLCs. (F) Mean intensity plot by RPM of SUZ12 ChIP-seq intensity at PSPC1/TET1/SUZ12 common peak regions (within ± 5K bp around SUZ12 peak center, identified in ESCs) in D2 EpiLCs. (G) SUZ12 ChIP-seq tracks at the promoters of bivalent genes (*Fgf5, Nefl, Sall2*, and *Ror2*) in D2 EpiLCs. The numbers above each track indicate the maximum reads per million (RPM) value.

Next, as PRC2 is the key player in regulating bivalent genes, we addressed how the PSPC1-TET1 partnership and the PSPC1-*Neat1* opposing functions may impose upon PRC2 in regulating bivalent genes during the pluripotent state transition. We performed SUZ12 ChIP-seq analysis in the D2 EpiLCs of *Pspc1*WT/KO or *Neat1*WT/KO genotypes and control- or dTAG13-treated *Tet1-*degron cells (Figure 5C). We chose D2 EpiLCs because a high anti-correlation was observed between the *Pspc1*KO and *Neat1*KO RNA-seq data (Figure 4C) and D2 EpiLCs represent the formative state of pluripotency (Smith, 2017) where super-bivalency was established (Wang et al., 2021; Xiang et al., 2020). As expected, PRC2 chromatin binding intensity at SUZ12 peak regions (identified in ESCs) significantly decreases in D2 EpiLCs compared to ESCs (Figure 5D-E). Plotting the SUZ12 binding intensity at SUZ12 peaks from *Pspc1*KO, *Neat1*KO, and dTAG13-treated *Tet1*-degron D2 EpiLCs, we found that SUZ12 binding (measure by the mean intensity in RPM) decreases upon depletion of PSPC1 or TET1, but not *Neat1*, at both all-SUZ12 peak regions (Figure S5D) and the PSPC1/SUZ12/TET1 common peak regions (Figure 5F). Importantly, we observed more considerable reductions of SUZ12 binding at PSPC1/SUZ12/ TET1 common peaks than at all-SUZ12 peaks upon depletion of PSPC1 or TET1 (compare the Δ[mean intensity] in Figure 5F with Figure S5D).

These results suggest that both PSPC1 and TET1, but not *Neat1*, are required for PRC2 chromatin occupancy, reinforcing the observed physical and functional partnership between PSPC1 and TET1 in regulating bivalent promoters (e.g., *Fgf5, Nefl, Sall2*) during the ESC-to-EpiLC transition (Figure 5G).

### PSPC1 and TET1 act through *Neat1* to modulate PRC2 binding to bivalent gene transcripts and control stem cell bivalency

While a physical association between the PSPC1-TET1 partnership and PRC2 is highly speculated (Figure 2C-D) for the observed functional interactions among these factors, neither a previously published work (Wu et al., 2011) nor our current study (Figure 1 and data not shown) can detect the TET1 and PRC2 interaction or the interactions between PSPC1 and PRC2 subunits using a regular nucleosome-free co-IP protocol (Figure S6A, refer to Methods for details). However, using a nucleosome-containing co-IP protocol with micrococcal nuclease (MN) digestion of chromatin, we and others readily detected the physical associations between TET1 and PRC2 (Neri et al., 2013) and between PSPC1 and PRC2 subunit SUZ12 (Figure 6A), respectively, in ESCs, raising the possibility of nucleosomal DNA/RNA molecules for tethering the protein interactions. By examining the datasets of PRC2 subunit EZH2 PAR-CLIP-seq (photoactivatable ribonucleoside-enhanced crosslinking and immunoprecipitation followed by sequencing) (Kaneko et al., 2013) and our PSPC1 CLIP-seq (Guallar et al., 2018) in ESCs, we observed that both EZH2 and PSPC1 are enriched at the *Neat1* transcripts (Figure S6B). In addition, CLIP-qPCR analysis of PSPC1 and EZH2 further confirmed the binding of both proteins to *Neat1* transcripts in WT ESCs (Figure 6B). These results support a tethering function of *Neat1* in mediating the PSPC1-PRC2 interaction. Furthermore, consistent with EZH2 direct binding to RNAs (Long et al., 2020), we found that EZH2 binding affinity to *Neat1* is not affected upon loss of *Pspc1* or *Tet1* in ESCs (Figure 6B) or EpiLCs (Figure S6D), suggesting that PRC2 binding to *Neat1* is independent of other RBPs such as PSPC1 irrespective of pluripotency states.

**Figure 6.**
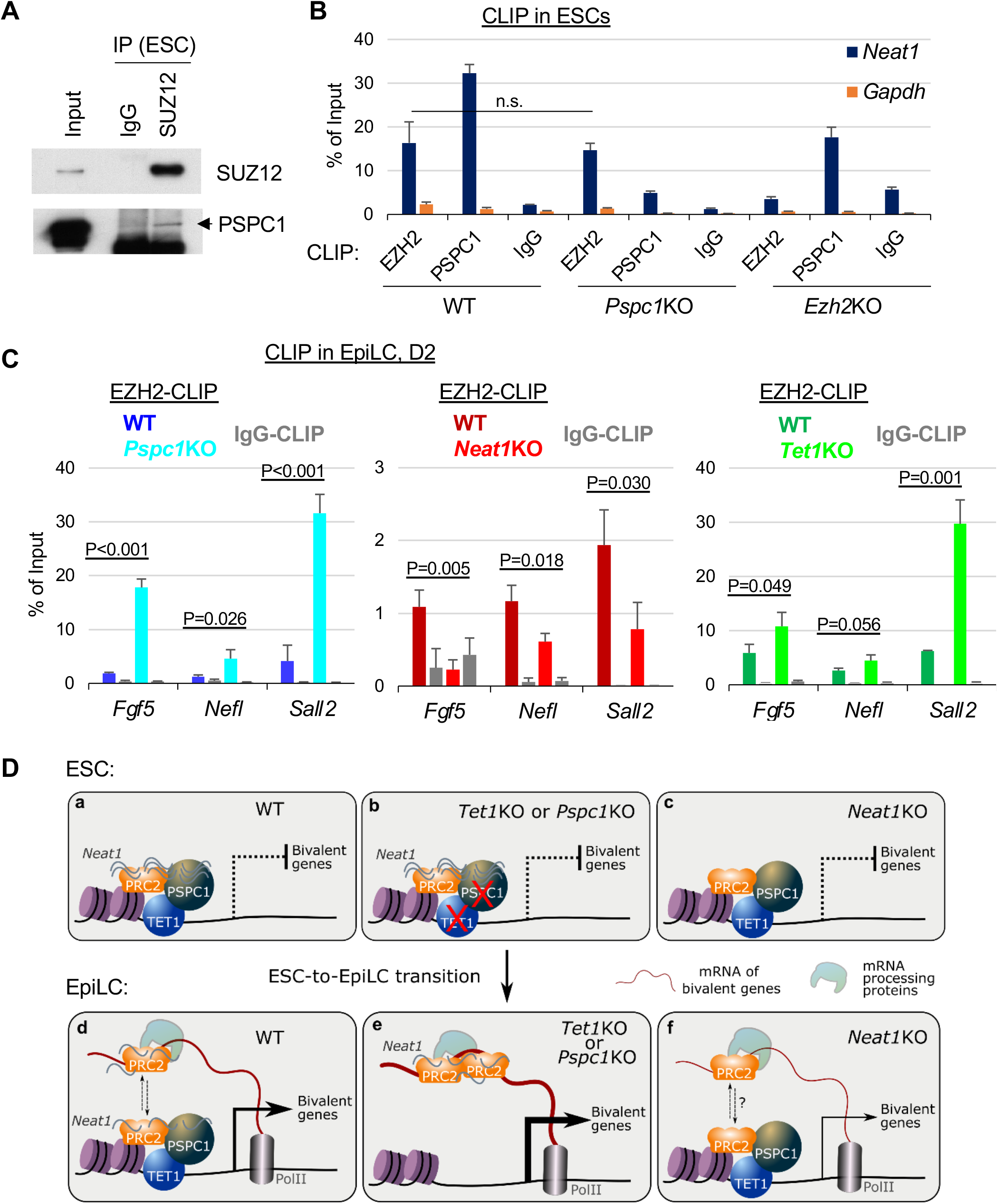
PSPC1, TET1, and *Neat1* modulate PRC2 affinity to nascent bivalent gene transcripts during bivalent gene activation. (A) Co-IP of PSPC1 and SUZ12 in ESCs using a nucleosome-containing protocol (see Methods in detail). (B) EZH2 and PSPC1 CLIP-qPCR analysis of *Neat1* in WT, *Pspc1*KO, and *Ezh2*KO ESCs. *Gapdh* serves as a negative control. P-value is from two-tailed T-test, and “n.s.” denotes statistically non-significant. Error bars represent the standard deviation of triplicates. (C) EZH2 CLIP-qPCR analysis of bivalent gene mRNAs (*Fgf5, Nefl, Sall2*) in D2 EpiLCs of different genotypes (WT vs. KO). P-value is from the two-tailed T-test. Error bars represent the standard deviation of triplicates. (D) The working model. In ESCs (WT), *Neat1* (short isoform, *Neat1_1*) tethers the chromatin-bound proteins TET1, PSPC1, and PRC2 at bivalent gene promoters (Panel a). Bivalent genes are not activated in either *Tet1*KO/*Pspc1*KO (Panel b) or *Neat1*KO (Panel c) ESCs. When bivalent genes are activated during pluripotent state transition (accompanied by downregulation of *Neat1_1*, with no expression of *Neat1_2* yet), nascent mRNA acts as a decoy to evict PRC2 from chromatin. In EpiLCs (WT), a dynamic balance is maintained between PRC2 chromatin occupancy and RNA binding (shown in up/down arrows) finetuning the expression of bivalent genes (Panel d). In *Tet1*KO or *Pspc1*KO EpiLCs, more PRC2 proteins bind to mRNAs and are displaced or evicted from chromatin, inducing enhanced bivalent gene transcription (Panel e). Without *Neat1* (during the pluripotent state transition in WT cells or upon genetic KO), PRC2 binding affinity to mRNAs (and possibly mRNA-processing-associated proteins) is compromised, which causes reduced bivalent gene activation (Panel f).

Since PRC2 has a higher affinity to RNA than DNA or histone, the nascent mRNAs during transcription activation decoy PRC2 and promote PRC2 eviction from chromatin (Wang et al., 2017a; Wang et al., 2017b). Whether and how the TET1-PSPC1-*Neat1* molecular interplay may modulate PRC2 affinity to nascent bivalent gene transcripts are particularly relevant in further unraveling the molecular mechanisms underlying stem cell bivalency. To address this, we performed EZH2 CLIP-qPCR analysis at the same D2 EpiLCs of *Pspc1* WT/KO, *Neat1* WT/KO, and *Tet1* WT/KO genotypes. We first confirmed that EZH2 protein levels are not affected upon loss of *Pspc1, Neat1*, or *Tet1* in ESCs and D2 EpiLCs (Figure S6C). We then compared EZH2 affinity to the transcripts of bivalent genes (e.g., *Egf5, Nefl*, and *Sall2*) activated during the ESC-to-EpiLC transition (Figure 3E-F). We found that EZH2 binding affinity to these mRNA transcripts increases upon the loss of *Pspc1* or *Tet1* (Figure 6C), accompanied by decreased PRC2 chromatin binding at promoters (Figure 5G). However, EZH2 affinity to these mRNA transcripts decreases upon the loss of *Neat1* (Figure 6C). We also performed PSPC1 CLIP-qPCR in D2 EpiLCs and found that, interestingly, while PSPC1 still binds to *Neat1*, it no longer binds to bivalent gene transcripts in EpiLCs upon the loss of *Neat1* or *Tet1* (Figure S6E).

Taken together, our results reveal that TET1 and PSPC1 inhibit the PRC2 affinity to bivalent mRNA transcripts, while *Neat1* facilitates PRC2 binding affinity to bivalent mRNAs during pluripotent state transition.

## DISCUSSION

In this study, we establish a stem cell paradigm whereby PSPC1 and TET1 prevent transcriptional activation of bivalent genes in ESCs and finetune the bivalent gene transcription during the ESC-to-EpiLC transition by promoting PRC2 chromatin occupancy and restricting the PRC2 affinity to the bivalent gene transcripts, respectively, partly through *Neat1*-mediated tethering of PRC2 to the chromatin (Figure 6Da,d). Upon the loss of *Tet1* or *Pspc1, Neat1* maintains its expression and positively mediates transcriptional activation of bivalent genes, likely through promoting PRC2 binding to the nascent mRNAs (Figure 6De). Our study thus provides mechanistic insights into how a dynamic balance between PRC2 chromatin occupancy and scanning of mRNA is maintained during the ESC-to-EpiLC transition (indicated by the up/down dashed arrows of Figure 6Dd). Without *Neat1* (i.e., *Neat1* KO), the balance of PRC2 chromatin occupancy and RNA binding affinity may be altered in favor of the former, resulting in attenuated bivalent gene transcription (Figure 6Df). While *Neat1* function in modulating PRC2 chromatin occupancy was reported (Wang et al., 2019), its function in promoting PRC2 binding affinity to bivalent gene transcripts when PSPC1 and/or TET1 are downregulated is an unexpected finding. In recent years, phase separation in the regulation of gene transcription has become an area of intense research (Hnisz et al., 2017). RNA Pol II acts in gene transcription through phase separation (Lu et al., 2018), and *Neat1* also scaffolds protein interactions of many RBPs that align to form paraspeckles by phase separation (Yamazaki et al., 2018). In our model, *Neat1* may facilitate phase separation of other mRNA processing proteins (i.e., ribonucleoprotein complex) for maintaining gene transcription and mRNA processing. This concept is supported by a recent proteomics study revealing that RNase treatment or Pol II inhibition reduces the chromatin fraction of RNA processing proteins (e.g., SF3AD, HNRNPU) while increasing the chromatin fraction of transcription factors and chromatin modifiers (Skalska et al., 2021). The nascent mRNA and other noncoding RNAs, including *Neat1*, may contribute to a dynamic matrix or phase-separated compartments that regulate chromatin states and gene transcription (Creamer et al., 2021; Skalska et al., 2021).

While a published study establishes a catalytic activity-dependent role of TET1 in demethylating bivalent promoters (Figure S2B) for the super-bivalency in formative pluripotency (Xiang et al., 2020), our study delineates a catalytic activity-independent role of TET1 in preventing hyper-activation of bivalent genes and thus preserving the bivalency in ESCs and during the ESC-to-EpiLC transition. Our data also support the PRC2 “eviction” models (Wang et al., 2017a; Wang et al., 2017b) and provide detailed mechanistic insight into the proposed repressive role of TET1 during bivalent gene activation (Koh et al., 2011; Wu et al., 2011). The TET family of proteins (TET1/2/3) are dynamically expressed during embryonic development. TET1 and TET2 are expressed in ESCs, but only TET1 is expressed in EpiLCs and EpiSCs (Fidalgo et al., 2016). We previously identified that PSPC1 also interacts with TET2, and PSPC1 recruits TET2 to the RNA transcripts of endogenous retrovirus (ERV) elements for RNA demethylation (Guallar et al., 2018). In this study, we identified that PSPC1 independently interacts with TET1 and TET2 (Figure 1E-F) and contributes to bivalent gene repression through a TET1 catalytic activity-independent mechanism. It is important to note that both catalytic activity-dependent and - independent functions of TET1 significantly contribute to early embryonic development (Khoueiry et al., 2017; Koh et al., 2011). Therefore, we cannot exclude the potential role of PSPC1 in the catalytic activity-dependent function of TET1, especially in differentiation procedures other than the naïve-to-formative differentiation (Li et al., 2020). In the current study, we report a previously unexplored group of PSPC1 and TET1 co-binding genomic targets (i.e., *Pou5f1, Nanog*) that are actively transcribed without PRC2 occupancy (Figure 2E-F). We recently revealed that PSPC1 promotes Pol II engagement and activity for the actively transcribed genes by enhancing the phase separation and subsequent phosphorylation and release of polymerase condensates (Shao et al., 2021). A study in human MCF7 cancer cell line indicated that *Neat1* can also bind to active chromatin sites (West et al., 2014). Therefore, the molecular actions of PSPC1 (and its cognate LncRNA *Neat1*) on the actively transcribed genes and bivalent genes are likely different.

The lncRNA *Neat1* has two isoforms. The long isoform *Neat1_2* is essential for the assembly of paraspeckles (Jiang et al., 2017; Nakagawa et al., 2011). The short isoform *Neat1_1*, albeit also a paraspeckle component, plays various paraspeckle-independent roles (Fox et al., 2018; Li et al., 2017). Human (Chen and Carmichael, 2009) or mouse (Figure S1C) ESCs have no paraspeckle assembly and a low expression level of *Neat1_2* (Figure 4B). In our study, while *Neat1_1* expression decreases during ESC-to-EpiLC differentiation, *Neat1_2* is not yet expressed in D2 or D4 EpiLCs (Figure 4B). Therefore, downregulation of *Neat1_1* may be necessary for the proper differentiation of ESCs. During differentiation of human ESC (hESC), paraspeckles start to form with the expression of *Neat1_2* (Modic et al., 2019). TDP43 post-transcriptionally regulates alternative polyadenylation (APA) of *Neat1* to produce the long isoform *Neat1_2* required for efficient early differentiation from hESCs (Grosch et al., 2020; Modic et al., 2019). It is interesting that hESCs also resemble the formative pluripotency, equivalent to mouse EpiLCs in gene expression patterns and lineage differentiation potentials (Smith, 2017). Therefore, expression of *Neat1_1* while lacking paraspeckle assembly is conserved in both mouse and human pluripotency, akin to the conservation of stem cell bivalency in both mouse and human.

### Limitations of study

Our current study does not have direct evidence that the nascent mRNAs of bivalent genes are subjected to PRC2 dynamic binding during ESC-to-EpiLC transition, which requires CLIP-seq analysis of PRC2 in a combination of global run-on sequencing (GRO-seq) to measure the association of PRC2 with the nascent mRNAs. However, this does not compromise our proposed model, emphasizing differential chromatin occupancy versus RNA binding of PRC2 as critical for bivalent gene regulation. Furthermore, as discussed, we reported in another study that nascent RNAs could synergize with PSPC1 and promote Pol II activity by enhancing phase separation (Shao et al., 2021), although it remains to be determined whether *Neat1* competes or facilitates nascent RNAs in the polymerase condensates on bivalent genes. In addition, since *Neat1* expression decreases during the naive-to-formative transition, a high dose of *Neat1* may be repressive by tethering PRC2 at chromatin (Wang et al., 2019), whereas a low dose of *Neat1* (with PSPC1) may facilitate Pol II engagement and subsequent phosphorylation and release of polymerase condensates (Shao et al., 2021). This concept is consistent with the recent finding that low levels of (nascent) RNAs at regulatory elements promote polymerase condensate formation, whereas high levels of RNAs from active gene transcription can dissolve condensates (Henninger et al., 2021). Therefore, *Neat1* may have dosage-dependent effects on transcriptional regulation of bivalent genes through a possible phase separation mechanism. Finally, it was reported that TET proteins also have RNA binding capacity (He et al., 2021; He et al., 2016), the possibility that TET1 may directly function as an RBP in its partnership with PSPC1 cannot be discounted for transcriptional regulation of bivalent genes in pluripotent state transition and ESC differentiation.

## ACKNOWLEDGMENTS

We thank Dr. Wei Xie for providing the *Tet1* vectors for domain mapping, Dr. Fei Lan for providing the *Nono*KO ESCs, Dr. Taiping Chen for providing the *Dnmt1/3a/3b*TKO ESCs, and Dr. Francesco Neri for discussion on the TET1 co-IP protocol. This work in the Shen Laboratory was supported in part by the National Natural Science Foundation of China (31829003). Research in the Wang laboratory is funded by grants from the National Institutes of Health (R01GM129157, R01HD095938, R01HD097268, and R01HL146664) and by contracts from New York State Stem Cell Science (NYSTEM#C35583GG and C35584GG).

## AUTHOR CONTRIBUTIONS

X.H. conceived, designed, and conducted the study, performed bioinformatics analysis, and wrote the manuscript; N.B., Y.H, and D.G. performed experiments; Z.H., V.M., D.L., H.Z., and X.S. provided reagents and contributed to experiments; J.W. conceived the project, designed the experiments, and prepared and approved the manuscript.

## STAR METHODS

### KEY RESOURCES TABLE

| REAGENT or RESOURCE | SOURCE | IDENTIFIER |
| --- | --- | --- |
| **Antibodies** |  |  |
| TET1 | Millipore | Cat. 09-872 |
| TET1 | GeneTex | Cat. GTX125888 |
| PSPC1 | Santa Cruz | Cat. sc-84577 |
| PSPC1 | Bethyl | Cat. A303-206A |
| SUZ12 | Abcam | Cat. ab12073 |
| EZH2 | Cell Signaling | Cat. 5246S |
| SUZ12 | Abcam | Cat. ab12073 |
| V5 | Invitrogen | Cat. R960-25 |
| Mouse IgG | Millipore | Cat. 12-371 |
| Rabbit IgG | Millipore | Cat. PP64 |
| SIN3A | Abcam | Cat. ab3479 |
| PELP1 | Bethyl | Cat. A300-180A |
| TET2 | Abcam | Cat. ab124297 |
| NONO | Bethyl | Cat. A300-587A |
| SFPQ | Abcam | Cat. ab38148 |
| HA | Abcam | Cat. ab9110 |
| OCT4 | Santa Cruz | Cat. sc-5279 |
| NANOG | Bethyl | Cat. A300-397A |
| ACTIN | Sigma | Cat. A5441 |
| GAPDH | ProteinTech | Cat. 10494-1-AP |
| Histone3 | Abcam | Cat. ab1791 |
| VCL | Abcam | Cat. ab129002 |
| Mouse IgG HRP | Cell Signaling | Cat. 7076S |
| Rabbit IgG HRP | Jackson ImmunoRes | Cat. 715-175-151 |
| Trueblot Mouse IgG HRP | Rockland | Cat. RK-18-8817-31 |
| Trueblot Rabbit IgG HRP | Rockland | Cat. RK-18-8816-31 |
| DNA 5mC | Cell Signaling | Cat. 28692 |
| DNA 5hmC | Active Motif | Cat. 39769 |
| Anti-dsDNA | Abcam | Cat. ab27156 |
| **Cell lines** |  |  |
| Mouse ESC CCE | This study | N/A |
| *Pspc1* KO ESC | This study | N/A |
| *Tet1*-degron ESC | This study | N/A |
| Mouse ESC 46C | This study | N/A |
| *Neat1* KO ESC | This study | N/A |
| *Ezh2* KO ESC | This study | N/A |
| Mouse ESC V6.5 | Laboratory of R. Jaenisch | N/A |
| *Tet1* KO ESCs | Laboratory of R. Jaenisch | N/A |
| *Tet1/2/3* KO ESCs | Laboratory of R. Jaenisch | N/A |
| *Dnmt1/3a/3b* KO ESC | Laboratory of T. Chen | N/A |
| *Nono* KO ESC | Laboratory of F. Lan | N/A |
| **Chemicals, Peptides, and Recombinant proteins** |  |  |
| DMEM | GIBCO | Cat. 11965-092 |
| Heat inactivated FBS | GIBCO | Cat. 35-011-CV |
| Penicillin-Streptomycin | GIBCO | Cat. 15140-122 |
| L-Glutamine | GIBCO | Cat. 25030-081 |
| MEM NEAA | GIBCO | Cat. 11140-050 |
| 2-Mercaptoethanol | Sigma | Cat. M6250 |
| Puromycin | Sigma | Cat. P9620-10ML |
| Hygromycin | Omega | Cat. HG-80 |
| N2 | GIBCO | Cat. 17502-048 |
| B27 | GIBCO | Cat. 17504-044 |
| DMEM/F-12 | GIBCO | Cat. 11-330-032 |
| Neurobasal | GIBCO | Cat. 21-103-049 |
| LIF | Lab prep | N/A |
| GSK3i (CHIR99021) | Sigma | Cat. SML1046-25MG |
| MEKi (PD0325901) | Selleckchem | Cat. S1036 |
| Recombinant Fgf2 | R&D System | Cat. 233-FB |
| Recombinant Activin A | R&D System | Cat. 338-AC |
| 13C615N4 L-arginine | Cambridge Isotope | Cat. CNLM-539-H |
| 13C615N2 L-lysine | Cambridge Isotope | Cat. CNLM-291-H |
| dTAG-13 | Tocris | Cat. 6605 |
| **Oligonucleotides** |  |  |
| See Table S4 | N/A | N/A |
| **Deposited data** |  |  |
| PSPC1 ChIP-seq in ESC | This study | NCBI GEO: GSE182443 |
| SUZ12 ChIP-seq in ESC | This study | NCBI GEO: GSE182443 |
| SUZ12 ChIP-seq in EpiLC upon *Pspc1* KO, *Neat1* KO, and *Tet1*-degron treatments | This study | NCBI GEO: GSE182443 |
| *Pspc1* WT/KO RNA-seq in ESC and EpiLC | This study | NCBI GEO: GSE182443 |
| *Neat1* WT/KO RNA-seq in ESC and EpiLC | This study | NCBI GEO: GSE182443 |
| **Other data** |  |  |
| TET1 ChIP-seq in ESC | Wu et al., 2011 | NCBI GEO: GSE48519 |
| H3K27me3 ChIP-seq in ESC | Cruz-Molina et al., 2017 | NCBI GEO: GSE89211 |
| H3K4me3 and H3K27ac in ESC | Hon et al., 2014 | NCBI GEO: GSE48519 |
| 5mC and 5hmC meDIP-seq in ESC | Xiong et al., 2016 | NCBI GEO: GSE57700 |
| **Software and Algorithms** |  |  |
| STAR | 2.7.6a | https://github.com/alexdobin/STAR |
| Cufflinks | 2.2.1 | http://cole-trapnell-lab.github.io/cufflinks/ |
| Bowtie2 | 2.3.5 | http://bowtie-bio.sourceforge.net/bowtie2/ |
| IGV | 2.10.2 | https://software.broadinstitute.org/software/igv |
| samtools | 1.10 | http://www.htslib.org/ |
| PICARD | 2.18.5 | https://broadinstitute.github.io/picard/ |
| HOMER | 4.11.1 | http://homer.ucsd.edu/homer/ |
| MACS2 | 2.2.7 | https://github.com/macs3-project/MACS |
| NGSplot | 2.61 | https://github.com/shenlab-sinai/ngsplot |

## LEAD CONTACT AND MATERIALS AVAILABILITY

Further information and requests for resources and reagents should be directed to and will be fulfilled by the Lead Contact, Jianlong Wang.

## DATA AND CODE AVAILABILITY

ChIP-seq and RNA-seq data have been deposited in the Gene Expression Omnibus (GEO) with accession code: GSE182443.

## EXPERIMENTAL MODEL AND SUBJECT DETAILS

### Cell culture and *in vitro* differentiation

If not specified, mouse embryonic stem cells (ESCs) were cultured on 0.1% gelatin-coated plates and in ES medium: DMEM medium supplemented with 15% fetal bovine serum (FBS), 1000 units/mL recombinant leukemia inhibitory factor (LIF), 0.1 mM 2-mercaptoethanol, 2 mM L-glutamine, 0.1 mM MEM non-essential amino acids (NEAA), 1% nucleoside mix (100X stock), and 50 U/mL Penicillin/Streptomycin. The *Ezh2* KO ESCs (Shen et al., 2008) were cultured on 0.1% gelatin-coated plates and in naïve culture condition (2iL) using serum-free N2B27 medium (DMEM/F12 and Neurobasal medium mixed at a ratio of 1:1, 1x B27 supplement, 1x N2 supplement, 2 mM L-glutamine, 0.1 mM 2-mercaptoethanol, and 50 U/mL Penicillin/Streptomycin) supplemented with Gsk3β inhibitor (CHIR99021, 3 μM), Mek inhibitor (PD0325901, 1 μM), and LIF (1000 units/mL).

For SILAC labeling, ESCs were cultured in either SILAC heavy or light medium: ES medium with complete supplements but deficient in both L-lysine and L-arginine, and then supplemented with L-lysine and L-arginine (SILAC light) or ^13^C_6_^15^N_4_ L-arginine and ^13^C_6_^15^N_2_ L-lysine (SILAC heavy) amino acids (Cambridge Isotope Laboratories).

For *in vitro* ESC-to-EpiLC differentiation, ESCs were seeded on fibronectin-coated (10 μg/mL/cm^2^) plates and in ES medium. On the next day, the medium was switch to formative culture condition using serum-free N2B27 medium supplemented with Fgf2 (12 ng/mL) and Activin A (20 ng/mL) (FA).

### Affinity purification followed by mass spectrometry (AP-MS) analysis

We employed a previously validated ESC clone with the ectopic expression of the 3xFLAG tagged mouse *Tet1* (FL-*Tet1*) gene (Ding et al., 2015). Before the AP-MS experiment, the empty vector (EV)- and FL-Tet1-transfected ESCs were cultured in both SILAC heavy and light medium for 2 weeks with reciprocal labeling: Replicate#1, light of FL-Tet1 versus heavy of EV; Replicate#2, light of EV versus heavy of FL-Tet1. AP-MS was performed using our well-established protocols (Ding et al., 2015; Guallar et al., 2018; Huang et al., 2021). Briefly, the cell pellets were resuspended in ice-cold hypotonic buffer A (10 mM HEPES, pH 7.9, 1.5 mM MgCl_2_, 10 mM KCl, 0.5 mM DTT, 0.2 mM PMSF, and protease inhibitor cocktail (PIC, Sigma, P8340)) and incubated for 10 min on ice. The sample was centrifuged at 3,000 ×g for 5 min at 4°C, and the pellet containing nuclei was washed by resuspending with ice-cold buffer A and centrifuging at 10,000 x g for 20 min at 4°C. Then, nuclei were resuspended with ice-cold nuclear extract buffer C (20 mM HEPES, pH 7.9, 20% glycerol (v/v), 0.42 M NaCl, 1.5 mM MgCl_2_, 0.2 mM EDTA, 0.5 mM DTT, 0.2 mM PMSF, and PIC) and incubated at 4°C for 30 min with continuous mixing. Insoluble materials were pelleted by centrifugation at 25,000 x g for 20 min at 4°C. The supernatant was collected as nuclear extract (NE) and dialyzed against buffer D (20 mM HEPES, pH 7.9, 20% glycerol (v/v), 100 mM KCl, 0.2 mM EDTA, 0.5 mM DTT, 0.2 mM PMSF) for 3 h at 4°C. Then, 0.1 ml of Protein G agarose (Roche Diagnostic) equilibrated in buffer D containing 0.02% NP40 (buffer D-NP) was added to nuclear extracts in 15 ml tubes, in the presence of Benzonase (25 U/mL, Millipore 70664), and incubated/pre-cleared for 1 h at 4°C with continuous mixing. Precleared NE samples were incubated with pre-equilibrated anti-FLAG M2 affinity gel (Sigma, F2426) for 4 h at 4°C with continuous mixing. Five washes were performed with buffer D-NP. Bound material was eluted by incubation with buffer D-NP supplemented with 0.5 mg/mL 3xFLAG peptides (Sigma, F4799) for 2 h at 4°C with continuous mixing. The eluted proteins were concentrated with Amicon Ultra Centrifugal Filters (Millipore, UFC500396), boiled 5 min in Laemmli buffer, and fractionated on a 10% SDS-PAGE gel. The gel lanes were cut horizontally into pieces, and each was subjected to LC-MS/MS analysis (Huang et al., 2021). MS data were processed using MaxQuant software (Tyanova et al., 2016) with Andromeda search engine (Cox et al., 2011) against the International Protein Index (IPI) mouse protein sequence database (v.3.68). Carbamidomethylation (CAM) was set as the fixed modification, and methionine oxidation was set as the variable modification. Other parameters followed the default settings of MaxQuant. Then the protein quantification heavy/light ratio was calculated and exported by MaxQuant. Quantification results of MS spectra were exported from the MaxQuant Viewer software.

### Co-immunoprecipitation (co-IP)

Co-IP in regular (nucleosome-free) conditions was performed as previously described (Ding et al., 2015). The nuclei were purified with buffer A followed the AP-MS protocol. Then nuclei were resuspended with ice-cold lysis buffer (50 mM HEPES, pH 7.9, 250 mM NaCl, 0.1% NP-40, 0.2 mM EDTA, 0.2 mM PMSF, and PIC) and incubated at 4°C for 30 min with continuous mixing. About 2% of input was saved, then NE was diluted with 40% volume (v/v = 5:2) of dilution buffer (20 mM HEPES, pH 7.9, 20% glycerol (v/v), 0.05% NP-40, 0.2 mM EDTA, 0.2 mM PMSF, and PIC) as the co-IP buffer with the NaCl concentration of 180 mM. For antibody IP, the antibody and the same amount of mouse or rabbit IgG as control were added to the co-IP buffer, incubated with protein lysates overnight at 4°C with continuous mixing. Then, protein lysates were incubated with protein G-Agarose beads (#11243233001, Roche) for 2 h at 4°C with continuous mixing. For FLAG-IP, NE was incubated with anti-FLAG M2 affinity gel (Sigma, F2426) overnight at 4 °C with continuous mixing. The immunoprecipitates were washed four times with co-IP buffer (lysis buffer/dilution buffer = 5:2, v/v). Proteins were eluted from the beads by boiling in 1X SDS Laemmli loading buffer, followed by SDS-PAGE and western blot analysis.

Co-IP in nucleosome-containing conditions was performed following a published protocol (Neri et al., 2013). Briefly, cell pellets were resuspended in isotonic buffer (20 mM HEPES, pH 7.5, 100 mM NaCl, 250 mM Sucrose, 5 mM MgCl_2_, 5 μM ZnCl_2_, and PIC), incubated on ice for 5 min, and spun down 500 g for 5 min at 4°C. Then pellets were resuspended in isotonic buffer (no PIC) supplemented with 1% NP-40), vortexed for 10 seconds at the highest setting, incubated on ice for 5 min, and spun down 1000g for 5 min at 4°C. The pellets (nuclei) were resuspended in 200 μL digestion buffer (50 mM Tris-HCl, pH 8.0, 100 mM NaCl, 250 mM Sucrose, 0.5 mM MgCl_2_, 5 mM CaCl_2_, 5 μM ZnCl_2_, no PIC) and 1 uL of micrococcal nuclease (MNase, NEB, M0247S), incubated at 37°C water bath for 10 min. Then the MNase digestion was immediately stopped by adding 20 μL 0.5 M EDTA, and nuclei were spun down 13,000 g for 1 min at 4°C. The digested nuclei were resuspended in digestion buffer (with PIC), subjected to sonication with Bioruptor®, set 30 sec ON, 30 sec OFF, 5 cycles to break nuclei, and spun down 13,000 g for 5 min at 4°C. Protein supernatants were subjected to antibody incubation, washing with digestion buffer, protein elution, and SDS-PAGE, like the regular co-IP protocol.

The primary antibodies used for co-IP were: TET1 (Millipore, 09-872 and GeneTex, GTX125888), PSPC1 (Santa Cruz, sc-84577 and Bethyl, A303-206A), SUZ12 (Abcam, ab12073), EZH2 (Cell Signaling, 5246S), V5 (Invitrogen, R960-25), mouse IgG (Millipore, 12-371), and rabbit IgG (Millipore, PP64).

### Subcellular fractionation assay

The subcellular fractions of ESCs were extracted using the Subcellular Protein Fractionation Kit for Cultured Cells (Thermo, #78840). Briefly, about 5 × 10^6^ cells were used, and each subcellular fraction was collected following the standard protocol. Protein loadings were balanced according to the protein concentrations in the cytoplasmic fraction before western blot analysis.

### Gel filtration assay

Size exclusion chromatography (gel filtration assay) was performed as previously described (Ding et al., 2015). Briefly, nuclear extracts (10∼20 mg) of ESCs were applied to a gel filtration column (S400 HiPrep 16/60 Sephacryl, Amersham Biosciences), samples were eluted at 1 mL/min and continuously monitored with an online detector at a wavelength of 280 nm. Fractions were collected, concentrated, and subjected to western blot analysis with indicated antibodies.

### Domain mapping

The FLAG-tagged Tet1 full-length (FL) sequence and truncated variants in the PiggyBac expression vectors were obtained from previous studies (Costa et al., 2013; Zhang et al., 2016). The Pspc1 full-length sequence and truncated variants were PCR amplified and subcloned into the V5-tagged PiggyBac expression vectors. The TET1 and PSPC1 PiggyBac expression vectors and control empty vectors (EV) were transfected into ESCs with Lipofectamine 2000 Transfection Reagent (Invitrogen, 11668019) following the standard protocol. After drug selection, ESCs were expanded for co-IP. FLAG-IP (for TET1 FL and truncated variants) and V5-IP (for PSPC1 FL and truncated variants) were performed, followed by western blot analysis of PSPC1 and TET1, respectively.

### Western blot analysis

Western blot analysis was performed as previously described (Huang et al., 2017). Total proteins were extracted by RIPA buffer. Protein concentrations were measured by Bradford assay (Pierce, 23236), balanced, and subjected to SDS-PAGE analysis. The following primary antibodies were used: PSPC1 (Bethyl, A303-206A and Sigma, SAB4200503), TET1 (Millipore, 09-872 and GeneTex, GTX125888), SIN3A (Abcam, ab3479), PELP1 (Bethyl, A300-180A), TET2 (Abcam, ab124297), V5 (Invitrogen, R960-25), NONO (Bethyl, A300-587A), SFPQ (Abcam, ab38148), HA (Abcam, ab9110), OCT4 (Santa Cruz, sc-5279), NANOG (Bethyl, A300-397A), SUZ12 (Abcam, ab12073), EZH2 (Cell Signaling, 5246S), ACTIN (1:5000, Sigma, A5441), GAPDH (ProteinTech, 10494-1-AP), Histone 3 (H3) (Abcam, ab1791), and Vinculin (Abcam, ab129002).

### Immunofluorescence

Mouse embryonic fibroblasts (MEFs) and ESCs were grown on 24-well plates coated with 0.1% gelatin (w/v). After fixation with 4% paraformaldehyde (w/v) for 15 min, cells were permeabilized with 0.25% Triton X-100 (v/v) in PBS for 5 min and incubated with 10% BSA for 30 min at 37°C. For immunostaining, cells were incubated overnight at 4°C with PSPC1 antibody (Santa Cruz, sc-84577) in PBS with 3% BSA (w/v). The following day cells were incubated with fluorophore-labeled secondary antibodies for 1 h at RT. Cells were imaged with a Leica DMI 6000 inverted microscope.

### *Neat1* knockout (KO) ESCs

CRISPR/Cas9-mediated *Neat1*KO was performed as described in (Yin et al., 2015). Briefly, two vectors (with the same pGL3-U6-sgRNA-PGK-puromycin backbone, Addgene #51133) containing two sgRNA sequences (Table S4) targeting a 6K bp region containing the short isoform of Neat1 (*Neat1_1*) were cotransfected with a Cas9-expressing vector (pST1374-N-NLS-flag-linker-Cas9, Addgene #44758) into WT 46C ESCs by lipofectamine 2000 (Invitrogen). Transfected cells were selected with puromycin and blasticidin for 8 days before clones were picked. Then, individual ESC clones were expanded and subjected to genomic DNA extraction and PCR for genotyping screening. The KO clones were further confirmed by RT-qPCR analysis of *Neat1* expression.

### *Tet1*-degron knock-in (KI) and protein degradation

The CRISPR/Cas9 system was used to engineer ESCs for protein degradation of TET1 genetically. The 5’- and 3’-homology arms of *Tet1* were PCR amplified from genomic DNA. The P2A-2xHA-FKBP(F36V) fragment for N-terminal insertion and the mCherry and BFP sequences were PCR amplified from Addgene plasmids #91792, #104370, #104371, respectively. *Tet1* 5’- and 3’-homology arms, FKBP, and mCherry or BFP sequences were assembled by Gibson Assembly 2x Master Mix (NEB, E2611S) to obtain 5’arm-FKBP-BPF-3’arm and 5’arm-FKBP-mCherry-3’arm doner vectors in pJET1.2 vector (Thermo Scientific). CRISPR gRNA was subcloned into the pSpCas9(BB)-2A-Puro (PX459) vector (gRNA sequence in Table S4). ESCs were transfected with the two donor vectors and CRISPR vectors using Lipofectamine 2000 (Invitrogen). After two days of puromycin selection, double-positive cells were sorted out for mCherry and BFP and seeded on a 96-well plate with single-cell per well using the BD Influx Cell Sorter. Cells were expanded and genotyped by PCR, and protein degradation was confirmed by western blot analysis. Clones with a homozygous knock-in tag were further expanded and used for experiments.

The *Tet1*-degron ESCs were treated with either DMSO control or dTAG13 (500 nM in DMSO, Tocris, 6605) for rapid degradation of TET1 protein. ESCs were treated with dTAG13 for 2 days before differentiation, and then cells were treated with dTAG13 during the ESC-to-EpiLC differentiation. In the control group, cells were treated with DMSO in ESCs and during the ESC-to-EpiLC differentiation.

### Dot blot analysis

The genomic DNA dot-blot analysis of 5mC and 5hmC was performed following the DNA Dot Blot Protocol (Cell Signaling, #28692) with modifications. Briefly, genomic DNA of ESCs was extracted using Quick-DNA Miniprep Plus Kit (Zymo, D4068), and DNA concentration was measured by NanoDrop. Next, the same amount of DNA was denatured with 10X DNA denaturing buffer (1 M NaOH and 0.1 M EDTA) and incubated at 95°C for 10 min, which was then immediately mixed with an equal volume of 20X SSC buffer, pH 7.0 (Invitrogen, 15557044) and chilled on ice. The DNA samples were diluted with a pre-determined amount and loaded on the positive-charged Nelyon membrane (GE Amersham, RPN2020B) using a vacuum chamber (Minifold, SRC-96). The membrane was dried, auto-crosslinked with 1200 x100 μJ/cm^2^, and blocked with 5% milk/TBST for 1 h. Next, the membrane was incubated with 5mC (Cell Signaling, 28692) or 5hmC (Active Motif, 39769) antibodies, the same as the western blot analysis. Then, the membrane was stripped with the stripping buffer (Thermo Scientific, 21059) and reblotted with the dsDNA (Abcam, ab27156) antibody as the loading control.

### Chromatin immunoprecipitation (ChIP) and sequencing

ChIP assays were performed as previously described (Huang et al., 2017). Briefly, cell pellets were cross-linked with 1% (w/v) formaldehyde for 10 min at RT, followed by the addition of 125 mM glycine to stop the reaction. Next, chromatin extracts were sonicated into 200–500 bp with Bioruptor®, set 30 sec ON, 30 sec OFF, 30 cycles, and immunoprecipitated with the following primary antibodies: PSPC1 (Santa Cruz, sc-84577), SUZ12 (Active Motif, 39357), and rabbit IgG (Millipore, PP64) overnight at 4°C with continuous mixing. The immunoprecipitated DNA was purified with ChIP DNA Clean & Concentrator columns (Zymo Research) and analyzed by qPCR with Roche SYBR Green reagents and a LightCycler480 machine. Percentages of input recovery were calculated. ChIP-qPCR primer sequences are listed in Table S4.

For ChIP-seq, 10% of sonicated genomic DNA was used as ChIP input. Libraries were prepared using the NEBNext Ultra II DNA library prep kit and index primers sets (NEB, 7645S, E7335S) following the standard protocol. Sequencing was performed with the Illumina HiSeq 4000 Sequencer according to the manufacturer’s protocol. Libraries were sequenced as 150-bp paired-end reads.

### ChIP-seq data processing

ChIP-seq data of histone marks H3K4me3 and H3K27ac in ESCs were downloaded from GSE48519 (Hon et al., 2014), data of H3K27me3 in ESCs were downloaded from GSE89211 (Cruz-Molina et al., 2017), and data of TET1 in ESCs were download from GSE26832 (Wu et al., 2011). ChIP-seq reads of biological replicates were combined. Briefly, reads were pre-processed by trim_galore (v0.6.3) and aligned to the mm9 mouse genome using the bowtie2 (v2.3.4) program, and the parameters were “-X 1000 --no-mixed --no-discordant”. The aligned reads were exported (-F 0×04 -f 0×02) and sorted with samtools. Duplicates were removed with MarkDuplicates function in the PICARD (v2.14.0) package. All Bam files were converted to a binary tiled file (tdf) and visualized using IGV (v2.7.2) software.

All ChIP-seq peaks were determined by the MACS2 program (v.2.0.10). PSPC1 ChIP peaks in WT cells were called using the *Pspc1* KO ChIP-seq as the control data, and SUZ12 ChIP peaks were called using the input ChIP-seq as the control data and all other parameters were the default settings. All peaks were annotated using the annotatePeaks module in the HOMER program (v4.11) against the mm9 genome. A target gene of a called peak was defined as the nearest gene’s transcription start site (TSS) with a distance to TSS less than 50 kb. Heatmaps and mean intensity curves of ChIP-seq data at specific genomic regions were plotted by the NGSplot program (v2.61) centered by the middle point “(start+end)/2” of each region.

ChIP-seq correlation analysis of PSPC1 and other factors was performed with an in-house Python program as previously described (Ding et al., 2015). A phi correlation coefficient was used to calculate the correlation between the ChIP peaks of every two ChIP-seq data. Heatmap of correlations was shown with the Java TreeView (v1.1.6) program.

### 5mC and 5hmC DNA immunoprecipitation (DIP) data analysis

5mC and 5hmC DIP-seq data in ESCs were downloaded from GSE57700 (Xiong et al., 2016). Reads were aligned to the mouse genome mm9 using the bowtie (v1.0.0) program, with parameters -m 1 -v 2 --best --strata. The duplicated reads of the aligned data were removed, then filtered reads were sorted with samtools (v0.1.19). The reads per million (RPM) values of 5mC and 5hmC DIP-seq at each TET1 ChIP-seq peak region were calculated with the NGSplot program (v2.61) and shown in Boxplots using R. P-value was calculated from two-sided Mann-Whitney test.

### RT-qPCR

Total RNA was extracted using the GeneJet RNA Purification Kit (Thermo Scientific, K0732). Reverse transcription was performed, and cDNA was generated using the qScript kit (Quanta, 95048). Relative expression levels were determined using a QuantStudio 5 Real-Time PCR System (Applied Biosystems). Gene expression levels were normalized to *Gapdh*. Error bars indicate standard deviation for the average expression of three technical replicates. Primers for RT-qPCR are listed in Table S4.

### Cross-linking immunoprecipitation (CLIP) qPCR

UV cross-linking and immunoprecipitation (CLIP) were performed according to the eCLIP-seq protocol (Van Nostrand et al., 2016) with modifications. Briefly, cells in culture were washed with ice-cold PBS and cross-linked in PBS with UV type C (254 nm) at 400 mJ/cm^2^ on ice. Next, cells were scraped, pelleted, and lysed in CLIP lysis buffer (50 mM Tris-HCl, pH 7.4, 100 mM NaCl, 1% NP-40, 0.1% SDS, 0.5% sodium deoxycholate) supplemented with proteinase and RNase inhibitors, and incubated on ice for 1 h. The lysate was briefly sonicated with Bioruptor®, set 30 sec ON, 30 sec OFF, 5 cycles to break DNA. Next, Turbo DNase (2U/μL, 1:500, Invitrogen, AM2238) was added, and the lysate was incubated in a 37°C water bath for 15 min, followed by centrifuge 15,000 g for 15 min at 4°C. Primary antibodies PSPC1 (Bethyl, A303-206A) and EZH2 (Cell Signaling, 5246S) or rabbit IgG (Millipore, PP64) were incubated with Protein-G dynabeads (Invitrogen) for 1 h at room temperature. Then the lysate and the beads were mixed overnight at 4°C with rotation. The next day, the beads were washed with wash buffer (low salt, 20 mM Tris-HCl, pH 7.4, 10 mM MgCl_2_, 0.2% Tween-20) and high salt wash buffer (50 mM Tris-HCl, pH 7.4, 1 M NaCl), and digested with Proteins K to elute RNA. The input and CLIP RNAs were purified with the RNA Clean & Concentrator-5 kit (Zymo, R1015) followed by RT-qPCR analysis. Percentages of input recovery were calculated. CLIP-qPCR primer sequences are listed in Table S4.

### RNA-seq and data analysis

100 ng total RNA was processed for RNA-seq library construction using the Ovation Mouse RNA-seq kit (NuGEN, #0348–32) following the manufacturer’s protocol. Massively parallel sequencing was performed on an Illumina HiSeq 4000 Sequencing System. Libraries were sequenced as 150-bp paired-end reads. For RNA-seq data processing, reads were aligned to the mouse genome mm9 using STAR (v2.7.6a) with the default settings. Transcript assembly and differential expression analyses were performed using Cufflinks (v2.2.1). Assembly of novel transcripts was not allowed (-G). Other parameters of Cufflinks were the default setting. The summed FPKM (fragments per kilobase per million mapped reads) of transcripts sharing each gene_id was calculated and exported by the Cuffdiff program. In the gene expression matrix, a value of FPKM+1 was applied to minimize the effect of low-expression genes. P-values were calculated using a T-test. Differentially expressed genes (DEGs) were determined by two-sided T-test P-value<0.05 and fold-change>1.5. Boxplots for expression were generated using R. P-value was calculated from the two-sided Mann-Whitney test.

PCA-analysis was performed for RNA-seq data from different batches. Batch effects were adjusted by *ComBat* function implemented in the *sva* Bioconductor package (v.3.18.0). The expression data matrix was imported by Cluster 3.0 software for PCA analysis. PC values were visualized with the plot3d function in the rgl package from CRAN. All R scripts were processed on the R-Studio platform (v3.6.1).

### Gene ontology (GO) analysis

Gene ontology (GO) analyses were performed using the DAVID gene ontology functional annotation tool (https://david.ncifcrf.gov/tools.jsp) with all NCBI *Mus musculus* genes as a reference list.

