## Supplemental Figures&Legends for "A TET1-PSPC1-*Neat1* molecular axis modulates PRC2 functions in controlling stem cell bivalency"

### Supplementary Figure Legends

#### Figure S1. Identification of PSPC1 as a novel partner of TET1 in ESCs. Related to Figure 1.

(A) SILAC quantification of a PSPC1 peptide NLSPVVSNELLEQAFSQFGPVEK by mass spectrometry. Quantification is based on the intensity from two replicates with reciprocal heavy/light labeling of FLAG-IP (TET1) and Control-IP (empty vector) MS experiments.

(B) Co-IP of PSPC1 and paraspeckle proteins NONO, SFPQ in ESCs, detected by western blot analysis of the antibodies against those proteins. IgG-IP serves as the negative control.

(C) Microscopy immunostaining images of WT MEFs and ESCs for PSPC1 (red) and DAPI (blue). The while arrows indicate paraspeckles.

(D) PSPC1 knockout *(Pspc1KO*) (two independent clones, C4 and C9) does not affect OCT4 levels in ESCs. Vinculin serves as a loading control.

(E) Gel filtration assay for the co-fractionation of TET1, TET2, and PSPC1 in ESCs. Two potential protein complexes: Complex I (in the blue rectangle) containing TET1/TET2/PSPC1 and Complex II (in the red rectangle) containing TET2/PSPC1 are indicated.

(F) Domain mapping of *Tet1* variants that interact with wildtype PSPC1. FLAG-tagged full length (FL) and different variants of *Tet1* are indicated on top. Co-IP is performed with FLAG-IP of TET1 fragments followed by western blot analysis of PSPC1. The black arrows on the bottom show the correct size of expressed TET1 protein fragments. Empty vector (EV) serves as the negative control. DSBH denotes the double strain B helix domain of TET1.

(G) Domain mapping of *Pspc1* variants that interact with wildtype TET1. V5-tagged full length (FL) and different variants of *Pspc1* are indicated on top. Co-IP is performed with V5-IP of PSPC1 fragments followed by western blot analysis of TET1. Note that there is a nonspecific band of V5 at 35 KDa. The black arrows on the bottom indicate the correct size of expressed PSPC1 variants. DBHS (drosophila behavior/human splicing), RRM (RNA recognition motifs), and NOPS (NonA/paraspeckle) domains of PSPC1 are indicated. Empty vector (EV) serves as the negative control.

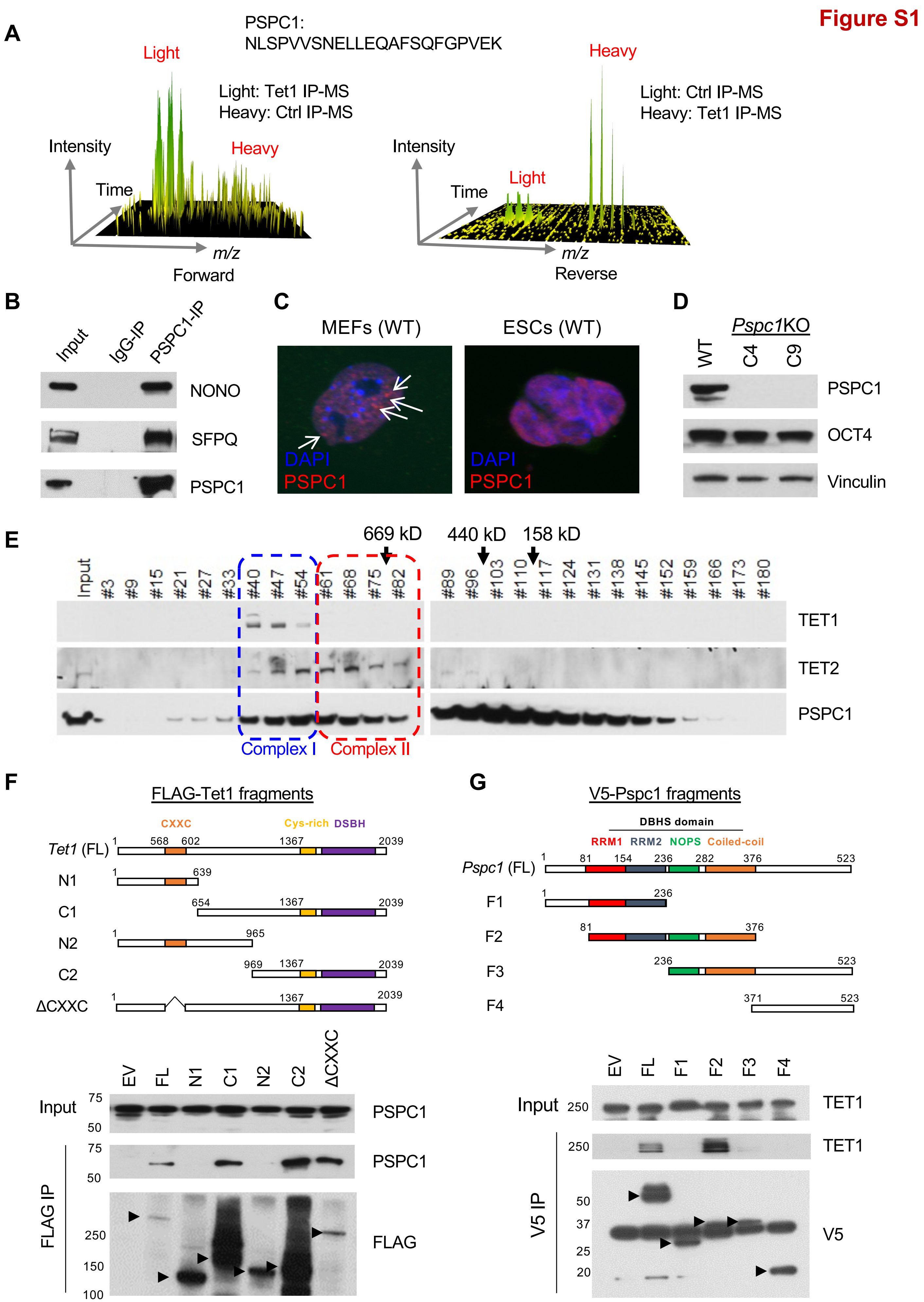

Figure S1
A
PSPC1:
NLSPVVSNELLEQAFSQFGPVEK
Light
Light: Tet1 IP-MS
Heavy: Ctrl IP-MS
Intensity
Heavy
Time
m/z
Forward
Heavy
Light: Ctrl IP-MS
Heavy: Tet1 IP-MS
Intensity
Light
Time
m/z
Reverse
B
Input
IgG-IP
PSPC1-IP
NONO
SFPQ
PSPC1
C
MEFs (WT)
ESCs (WT)
DAPI
PSPC1
DAPI
PSPC1
D
Pspc1KO
WT
C4
C9
PSPC1
OCT4
Vinculin
E
669 kD
440 kD
158 kD
Input
#3
#9
#15
#21
#27
#33
#40
#47
#54
#61
#68
#75
#82
#89
#96
#103
#110
#117
#124
#131
#138
#145
#152
#159
#166
#173
#180
TET1
TET2
PSPC1
Complex I
Complex II
F
FLAG-Tet1 fragments
CXXC
Cys-rich
DSBH
Tet1 (FL)
N1
C1
N2
C2
ΔCXXC
1
568
602
1367
2039
639
654
965
969
EV
FL
N1
C1
N2
C2
ΔCXXC
Input
75
50
PSPC1
PSPC1
FLAG IP
250
150
100
FLAG
G
V5-Pspc1 fragments
DBHS domain
RRM1
RRM2
NOPS
Coiled-coil
Pspc1 (FL)
F1
F2
F3
F4
1
81
154
236
282
376
523
371
EV
FL
F1
F2
F3
F4
Input
250
TET1
TET1
V5 IP
50
37
25
20
V5

**Figure S2. PSPC1, TET1, and PRC2 co-localize at the bivalent gene promoters in ESCs. Related to Figure 2.**

(A) Overlap of the PSPC1 and TET1 ChIP-seq peaks in ESCs.

(B) Boxplots depicting quantification of DNA 5hmC and 5mC intensity at TET1 peak regions with or without PSPC1 occupancy. P-value is from the Mann-Whitney test. DNA 5hmC and 5mC DIP-seq data in ESCs are curated from (Xiong et al., 2016).

(C) ChIP-seq correlation analysis based on identified peaks of pluripotency-related transcription factors and epigenetic regulators in ESCs. A blue rectangle indicates TET1 ChIP-seq peaks are associated with its interacting partner PSPC1 and PRC2 subunits EZH2 and SUZ12.

(D) Gene ontology (GO) analysis for the PSPC1/TET1/SUZ12 common target genes.

**Figure S2**

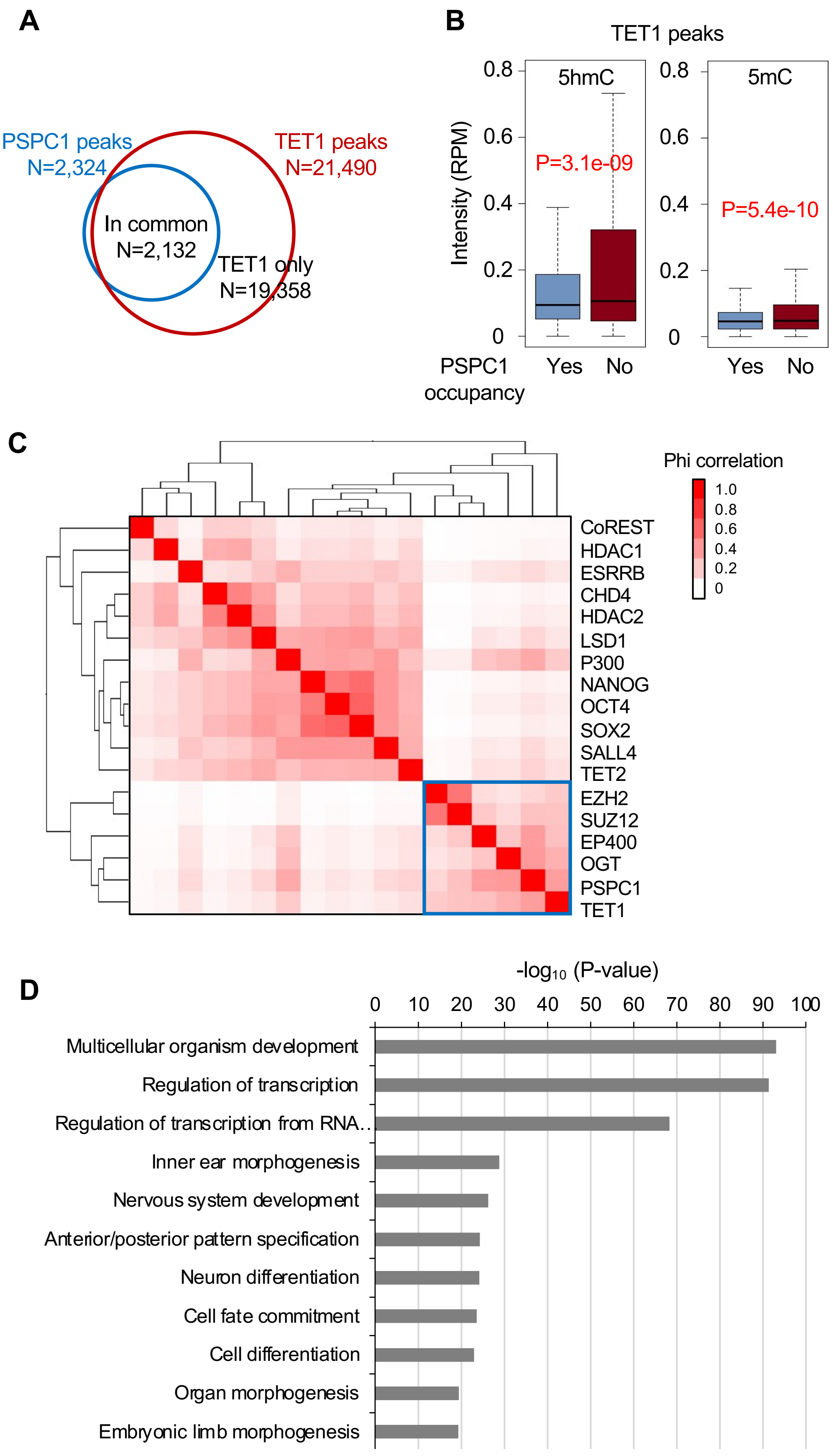
A
PSPC1 peaks
N=2,324
TET1 peaks
N=21,490
In common
N=2,132
TET1 only
N=19,358
B
TET1 peaks
5hmC
5mC
Intensity (RPM)
0.8
0.6
0.4
0.2
0
P=3.1e-09
P=5.4e-10
PSPC1 occupancy
Yes
No
C
Phi correlation
1.0
0.8
0.6
0.4
0.2
0
CoREST
HDAC1
ESRRB
CHD4
HDAC2
LSD1
P300
NANOG
OCT4
SOX2
SALL4
TET2
EZH2
SUZ12
EP400
OGT
PSPC1
TET1
D
-log10 (P-value)
0
10
20
30
40
50
60
70
80
90
100
Multicellular organism development
Regulation of transcription
Regulation of transcription from RNA..
Inner ear morphogenesis
Nervous system development
Anterior/posterior pattern specification
Neuron differentiation
Cell fate commitment
Cell differentiation
Organ morphogenesis
Embryonic limb morphogenesis

**Figure S3. PSPC1 and NONO negatively regulate bivalent gene activation in pluripotent state transition. Related to Figure 3.**

(A) Volcano plots depicting the differentially expressed genes (DEGs, P-value<0.05, fold-change>1.5) by comparing the WT and *Pspc1*KO cells at 3 time points (D0, D2, D4). Numbers of Down- and Up-regulated DEGs and some representative genes names are indicated.

(B) Pie charts depicting the overlap of the PSPC1-repressed (left) and -activated (right) DEGs in D2 and D4 EpiLCs. The numbers and percentages are indicated if these DEGs are also repressed (left) or activated (right) by PSPC1 in ESCs.

(C) Verification of the KO statuses of *Pspc1*KO and *Nono*KO ESCs by western blot analysis.

(D) RT-qPCR analysis of *Nanog* and lineage genes (*T, Eomes, Fgf8*) in WT and *Nono*KO ESCs during EpiLC differentiation.

Figure S3

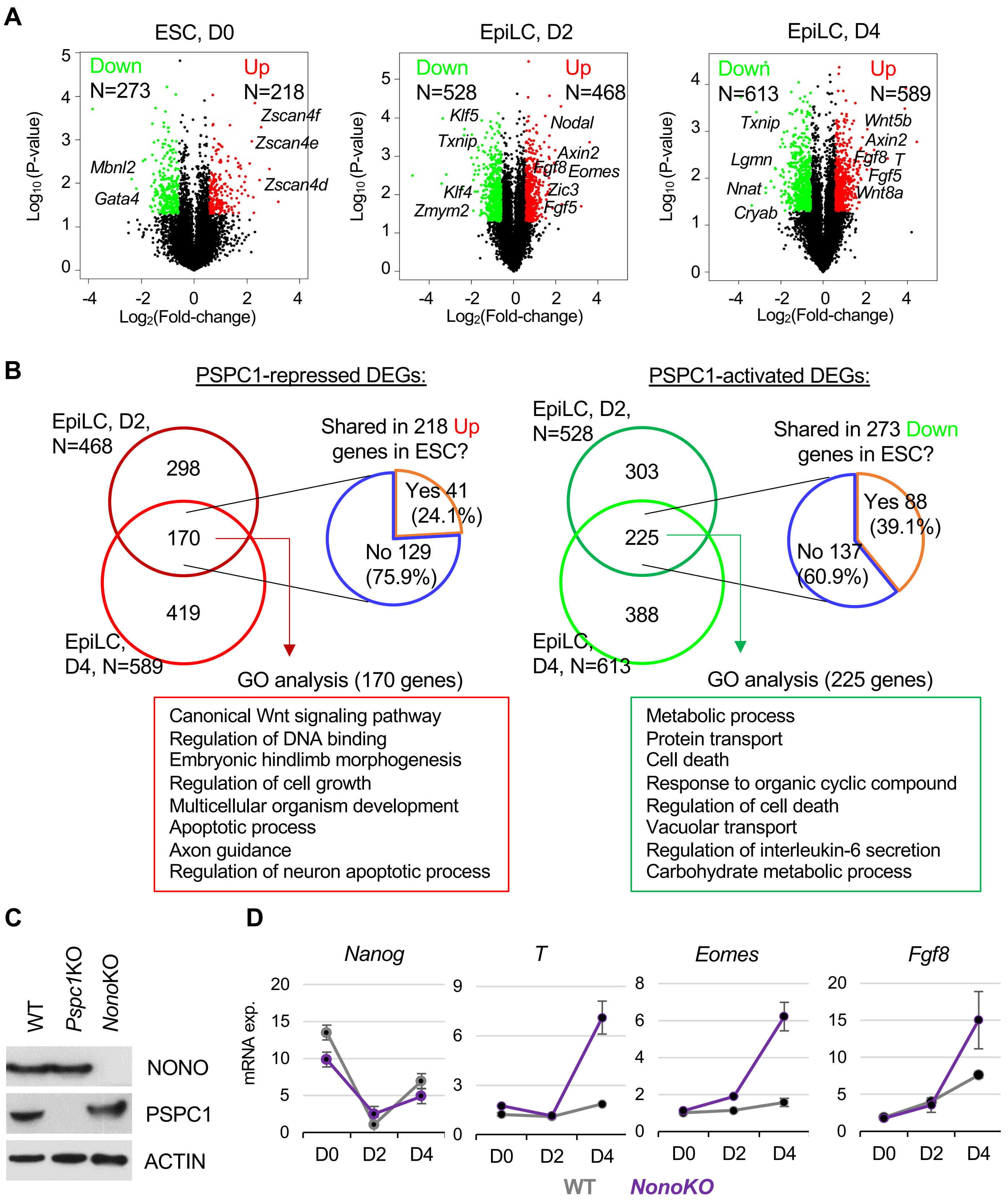

A
ESC, D0
EpiLC, D2
EpiLC, D4
Down
N=273
Up
N=218
Zscan4f
Zscan4e
Zscan4d
Mbnl2
Gata4
N=528
N=468
Klf5
Txnip
Nodal
Axin2
Fgf8
Eomes
Zic3
Fgf5
Klf4
Zmym2
N=613
N=589
Wnt5b
Lgmn
Nnat
Cryab
T
Wnt8a
Log10 (P-value)
Log2(Fold-change)
B
PSPC1-repressed DEGs:
PSPC1-activated DEGs:
EpiLC, D2, N=468
298
170
419
EpiLC, D4, N=589
Shared in 218 Up genes in ESC?
Yes 41 (24.1%)
No 129 (75.9%)
EpiLC, D2, N=528
303
225
388
EpiLC, D4, N=613
Shared in 273 Down genes in ESC?
Yes 88 (39.1%)
No 137 (60.9%)
GO analysis (170 genes)
Canonical Wnt signaling pathway
Regulation of DNA binding
Embryonic hindlimb morphogenesis
Regulation of cell growth
Multicellular organism development
Apoptotic process
Axon guidance
Regulation of neuron apoptotic process
GO analysis (225 genes)
Metabolic process
Protein transport
Cell death
Response to organic cyclic compound
Regulation of cell death
Vacuolar transport
Regulation of interleukin-6 secretion
Carbohydrate metabolic process
C
WT
Pspc1KO
NonoKO
NONO
PSPC1
ACTIN
D
Nanog
T
Eomes
Fgf8
mRNA exp.
D0
D2
D4
WT
NonoKO

**Figure S4. *Neat1* positively regulates activation of bivalent genes in pluripotent state transition. Related to Figure 4.**

(A) Volcano plots depicting the differentially expressed genes (DEGs, P-value<0.05, fold-change>1.5) by comparing the WT and *Neat1*KO cells at 3 time points (D0, D2, D4). Numbers of Down- and Up-regulated DEGs and some representative genes names are indicated.

(B) Scatter plots depicting the relative expression of all genes upon the loss of *Pspc1* (*Pspc1* KO/WT) or *Neat1* (*Neat1* KO/WT) at 3 time points (D0, D2, D4). The Pearson's product-moment correlation coefficient (*r*) and P-value of correlation are indicated in each plot.

(C) Scatter plots depicting the relative gene expression of DEGs (P-value<0.05, fold-change>1.5) upon the loss of *Pspc1* (*Pspc1* KO/WT) or *Neat1* (*Neat1* KO/WT) in ESCs (left) and D4 EpiLCs (right) from RNA-seq analysis. P-value is from the Fisher-extract test.

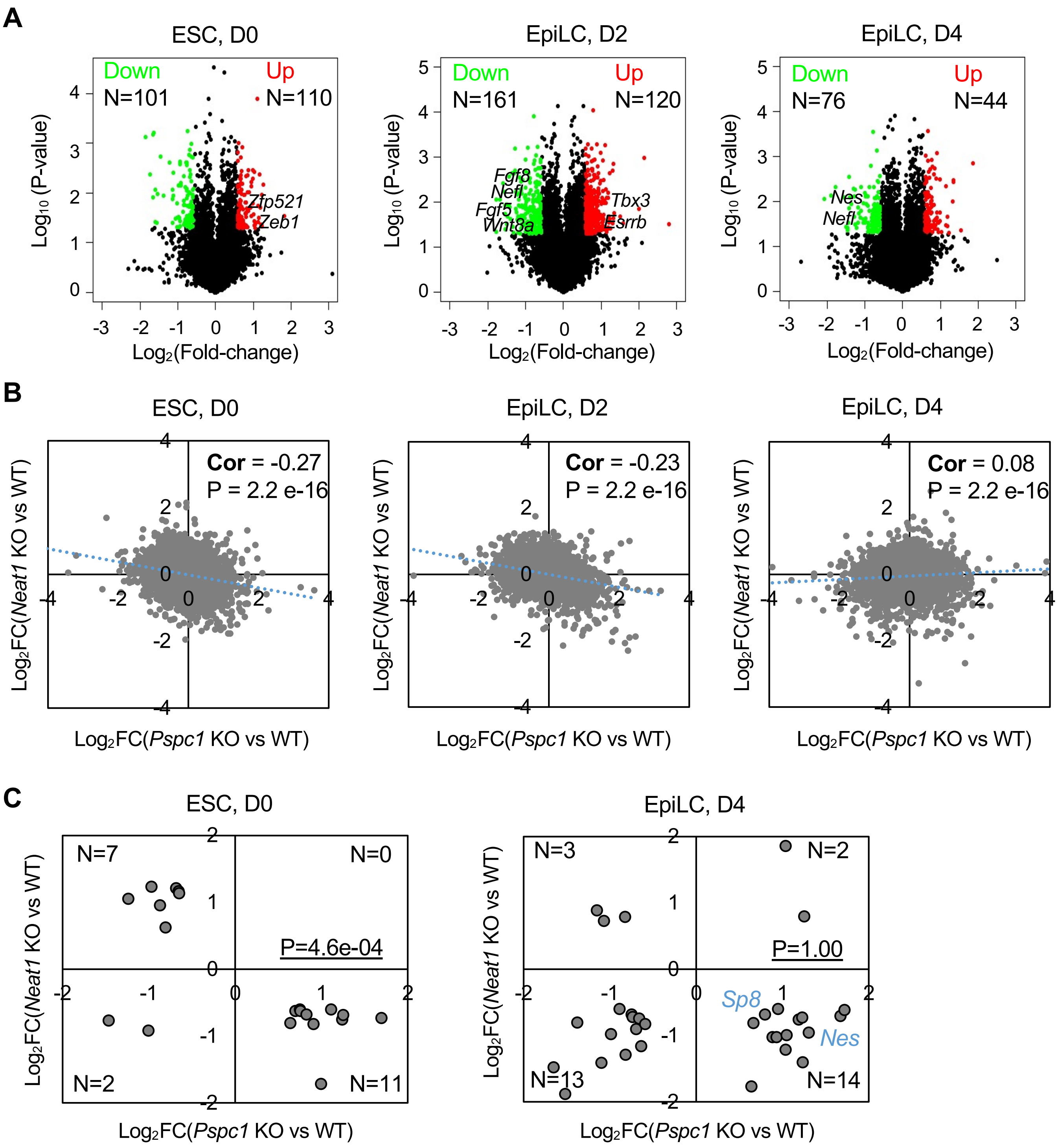
A
ESC, D0
Down
N=101
Up
N=110
Zfp521
Zeb1
EpiLC, D2
Down
N=161
Up
N=120
Fgf8
Nefl
Fgf5
Wnt8a
Tbx3
Esrrb
EpiLC, D4
Down
N=76
Up
N=44
Nes
Nefl
Log10 (P-value)
Log2(Fold-change)
B
ESC, D0
Cor = -0.27
P = 2.2 e-16
EpiLC, D2
Cor = -0.23
P = 2.2 e-16
EpiLC, D4
Cor = 0.08
P = 2.2 e-16
Log2FC(Neat1 KO vs WT)
Log2FC(Pspc1 KO vs WT)
C
ESC, D0
N=7
N=0
P=4.6e-04
N=2
N=11
EpiLC, D4
N=3
N=2
P=1.00
Sp8
Nes
N=13
N=14
Log2FC(Neat1 KO vs WT)
Log2FC(Pspc1 KO vs WT)

**Figure S5. Depletion of PSPC1 or TET1 accelerates PRC2 eviction from bivalent gene promoters. Related to Figure 5.**

(A) Western blot analysis of whole-cell lysate, cytoplasmic, and chromatin-bound fractions of PSPC1, TET1, and SUZ12 in WT and *Pspc1*KO (two independent clones, C4 and C9) ESCs. ACTIN and histone H3 are the loading controls of cytoplasmic and chromatin-bound fractions, respectively.

(B) SUZ12 ChIP-qPCR analysis at indicated gene promoters in WT and *Pspc1*KO ESCs. Error bars represent the standard deviation of triplicates.

(C) RT-qPCR analysis of the pluripotency gene (*Nanog*) and lineage genes (*T, Fgf5, Fgf8*) in *Tet1*-degron ESCs treated with control DMSO or dTAG-13 during EpiLC differentiation.

(D) Mean intensity plots (top) and heatmaps by reads per million (RPM) (bottom) depicting the SUZ12 ChIP-seq intensity at all SUZ12 peak regions (within ± 5K bp around SUZ12 peak center, identified in ESCs) in D2 EpiLCs of different genotypes (*Pspc1* WT/KO, *Neat1* WT/KO) or with other treatments (*Tet1*-degron with control/dTAG13).

Figure S5

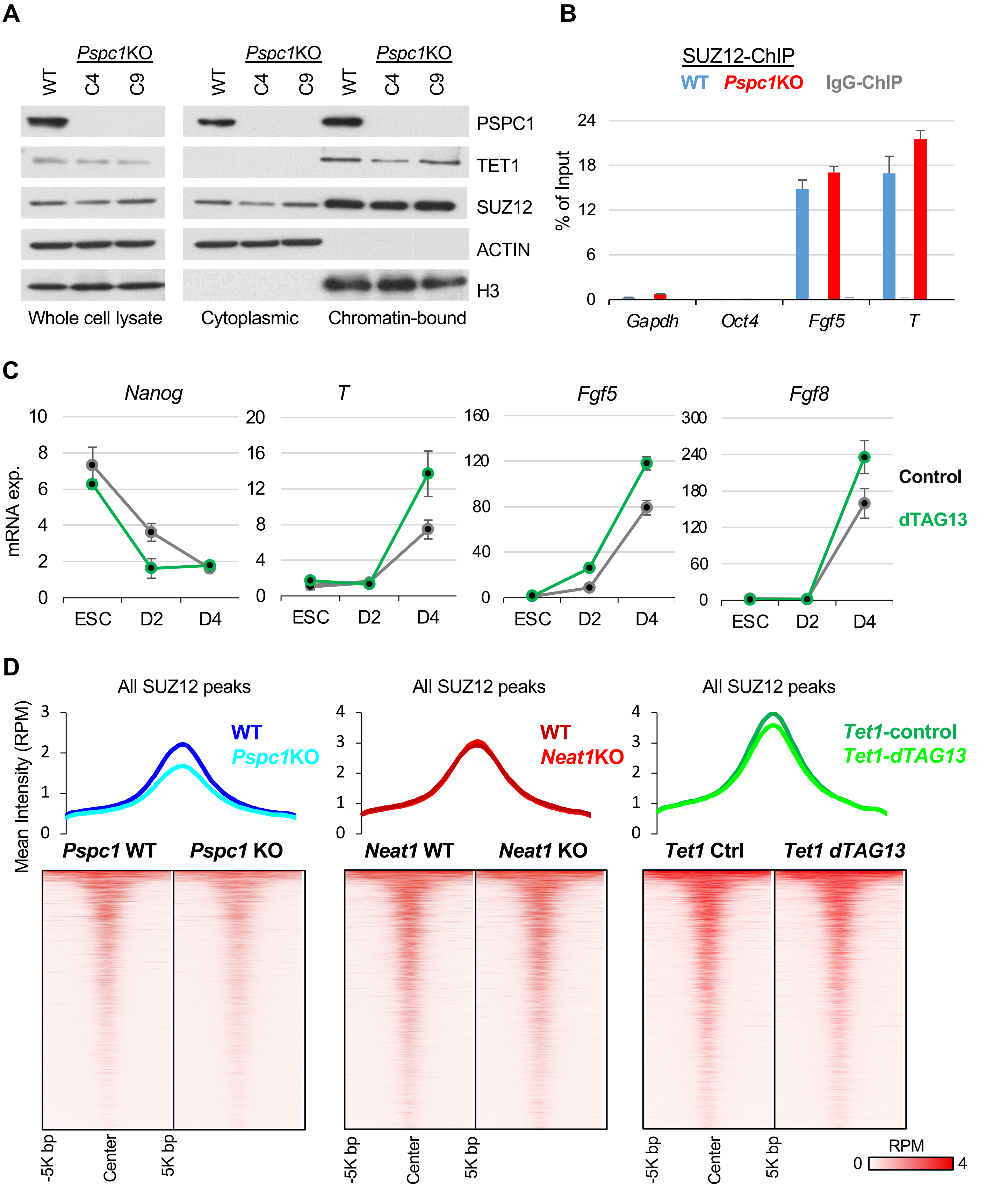
A
Pspc1KO
WT
C4
C9
PSPC1
TET1
SUZ12
ACTIN
H3
Whole cell lysate
Cytoplasmic
Chromatin-bound
B
SUZ12-ChIP
WT Pspc1KO IgG-ChIP
% of Input
Gapdh
Oct4
Fgf5
T
C
Nanog
T
Fgf5
Fgf8
mRNA exp.
ESC
D2
D4
Control
dTAG13
D
All SUZ12 peaks
Mean Intensity (RPM)
WT
Pspc1KO
Neat1KO
Tet1-control
Tet1-dTAG13
Pspc1 WT
Pspc1 KO
Neat1 WT
Neat1 KO
Tet1 Ctrl
Tet1 dTAG13
-5K bp
Center
5K bp
RPM
0
4

**Figure S6. PSPC1, TET1, and *Neat1* modulate PRC2 affinity to nascent bivalent gene transcripts during bivalent gene activation. Related to Figure 6.**

(A) The lack of physical association between PSPC1 and PRC2 under a nucleosome-free co-IP protocol. Co-IP of PSPS1 and PRC2 subunits EZH2 and SUZ12 was performed in ESCs using a nucleosome-free protocol (see details in Methods) followed by western blot analysis.

(B) Both EZH2 and PSPC1 binds to *Neat1* lncRNA. EZH2 PAR-CLIP-seq binding sites were processed from (Kaneko et al., 2013) and PSPC1 CLIP-seq was from (Guallar et al., 2018). The CLIP-seq tracks at *Neat1* locus in ESCs were indicated by green vertical lines (for EZH2) or peaks (for PSPC1).

(C) EZH2 total proteins are not changed by the loss of *Pspc1*, *Neat1*, or *Tet1*, or during the ESC-to-EpiLC transition, confirmed by western blot analysis with indicated antibodies. NANOG (a naïve marker) was notably downregulated in EpiLCs relative to ESCs, although its levels are relatively unaffected in KO relative WT ESCs or EpiLCs.

(D) EZH2 CLIP-qPCR analysis of *Neat1* in *Pspc1* WT/KO, *Neat1* WT/KO, and *Tet1* WT/KO D2 EpiLCs. P-value is from 2-tailed T-test, and “n.s.” denotes non-significant. Error bars represent the standard deviation of triplicates.

(C) PSPC1 CLIP-qPCR analysis of *Neat1* and bivalent gene mRNAs (*Fgf5, Neff, Sall2*) in D2 EpiLCs of different genotypes. Error bars represent the standard deviation of triplicates.

Figure S6

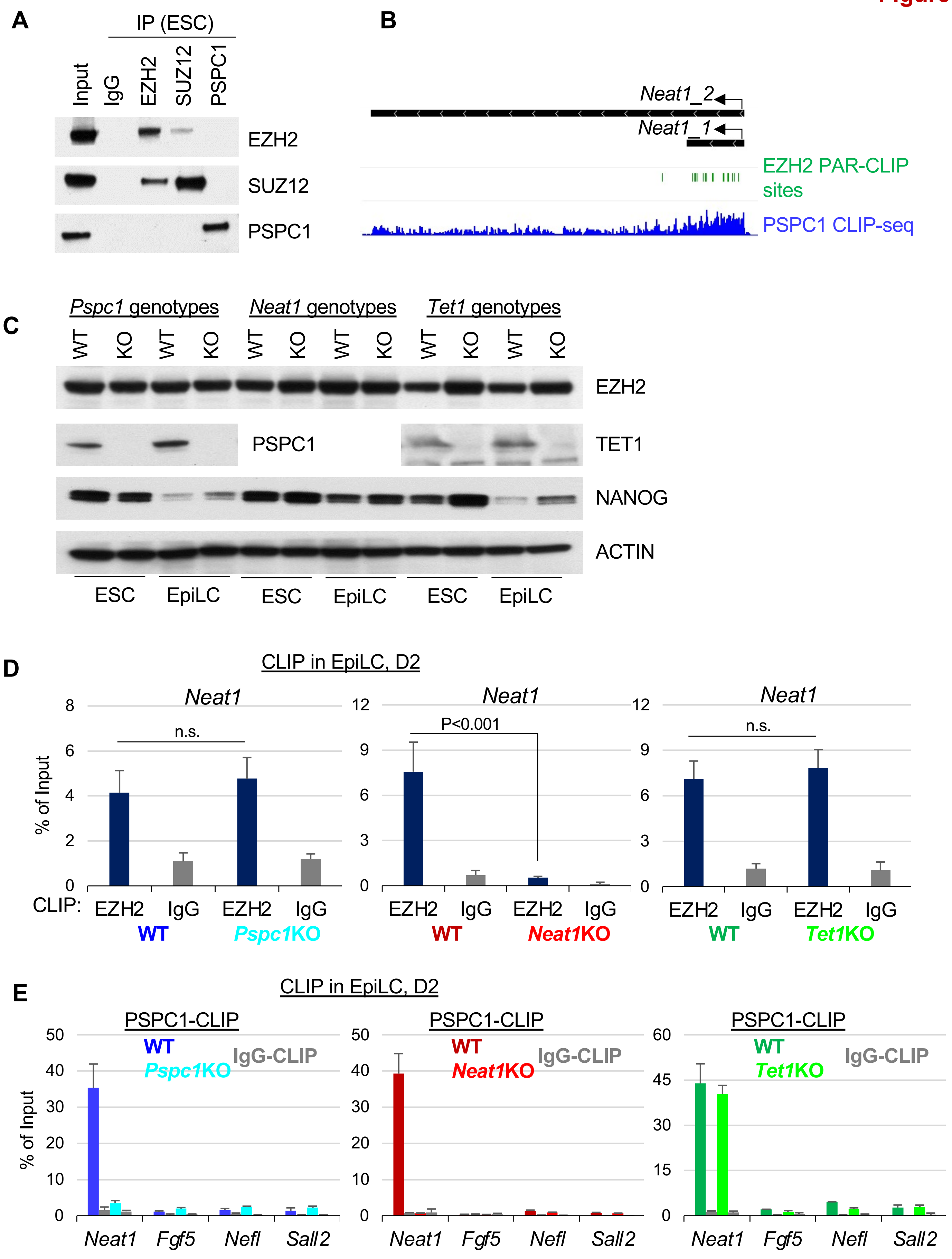

A
IP (ESC)
Input
IgG
EZH2
SUZ12
PSPC1
EZH2
SUZ12
PSPC1
B
Neat1_2
Neat1_1
EZH2 PAR-CLIP sites
PSPC1 CLIP-seq
C
Pspc1 genotypes
Neat1 genotypes
Tet1 genotypes
WT
KO
WT
KO
WT
KO
WT
KO
WT
KO
WT
KO
EZH2
PSPC1
TET1
NANOG
ACTIN
ESC
EpiLC
ESC
EpiLC
ESC
EpiLC
D
CLIP in EpiLC, D2
Neat1
Neat1
Neat1
% of Input
n.s.
P<0.001
n.s.
CLIP:
EZH2
IgG
EZH2
IgG
EZH2
IgG
EZH2
IgG
EZH2
IgG
EZH2
IgG
WT
Pspc1KO
WT
Neat1KO
WT
Tet1KO
E
CLIP in EpiLC, D2
PSPC1-CLIP
PSPC1-CLIP
PSPC1-CLIP
WT
Pspc1KO
IgG-CLIP
WT
Neat1KO
IgG-CLIP
WT
Tet1KO
IgG-CLIP
% of Input
Neat1
Fgf5
Nefl
Sall2
Neat1
Fgf5
Nefl
Sall2
Neat1
Fgf5
Nefl
Sall2
